# Narrowband multivariate source separation for semi-blind discovery of experiment contrasts

**DOI:** 10.1101/2020.03.09.983635

**Authors:** Marrit B. Zuure, Michael X Cohen

## Abstract

**Background:** Electrophysiological recordings contain mixtures of signals from distinct neural sources, impeding a straightforward interpretation of the sensor-level data. This mixing is particularly detrimental when distinct sources resonate in overlapping frequencies. Fortunately, the mixing is linear and instantaneous. Multivariate source separation methods may therefore successfully separate statistical sources, even with overlapping spatial distributions.

**New Method:** We demonstrate a feature-guided multivariate source separation method that is tuned to narrowband frequency content as well as binary condition differences. This method — comparison scanning generalized eigendecomposition, csGED — harnesses the covariance structure of multichannel data to find directions (i.e., eigenvectors) that maximally separate two subsets of data. To drive condition specificity and frequency specificity, our data subsets were taken from different task conditions and narrowband-filtered prior to applying GED.

**Results:** To validate the method, we simulated MEG data in two conditions with shared noise characteristics and unique signal. csGED outperformed the best sensor at reconstructing the ground truth signals, even in the presence of large amounts of noise. We next applied csGED to a published empirical MEG dataset on visual perception vs. imagery. csGED identified sources in alpha, beta, and gamma bands, and successfully separated distinct networks in the same frequency band.

**Comparison with Existing Method(s):** GED is a flexible feature-guided decomposition method that has previously successfully been applied. Our combined frequency- and condition-tuning is a novel adaptation that extends the power of GED in cognitive electrophysiology.

**Conclusions:** We demonstrate successful condition-specific source separation by applying csGED to simulated and empirical data.

## Introduction

EEG and MEG are prized for their high temporal precision, allowing us to link fast brain dynamics to cognition. However, their spatial precision is low compared to that of other measurement methods, with each sensor recording a linear and instantaneous mixture of populations of neurons from spatially distributed regions (Nunez & Srinivasan, 2006). Thus, many functionally diverse neural assemblies can contribute to the activity recorded from each MEG sensor. Fortunately, disparate neural sources generally produce unique electromagnetic fields, which have stable though spatially overlapping distributions at the scalp. Multichannel decomposition methods leverage these properties to attempt to isolate the component sources.

Many source separation approaches exist, and different approaches are designed to optimize different features (e.g., Beckmann & Smith, 2004; Brunton et al., 2016; Wang & Guo, 2019; Yang et al., 2004). We espouse the use of generalized eigendecomposition (GED), an advanced feature-guided multivariate source separation method. GED distinguishes itself from other methods by finding statistical sources (i.e., linear combinations of sensors) that maximally dissociate user-specified data of interest from reference data. This hypothesis-driven source separation provides improved flexibility and sensitivity over other multivariate source separation methods such as PCA and ICA.

Preparing multivariate data for GED requires the creation of two covariance matrices. One is constructed from data containing the features to be maximized (such as narrowband activity, a particular time window, or a subset of trials) and the other from data containing the features to be minimized (such as broadband activity, the entire trial, or another subset of trials). A GED applied to these covariance matrices returns a set of components, each consisting of an eigenvalue (indicating importance) and an eigenvector (a spatial filter, i.e., a set of sensor weights). These components capture sensor combinations that maximize the energy ratio between the covariance matrices. The eigenvectors can be used to construct component topographies and time series, the latter of which can be time-frequency decomposed for further inspection.

GED can identify frequencies of interest in the data by scanning the frequency spectrum for sources that maximally separate narrowband from broadband data (de Cheveigné & Arzounian, 2015; Nikulin et al., 2011). We adapted this technique to concurrently optimize for not only frequency content, but also condition differences. This choice was driven by the observation that experimental designs in neuroscience commonly harness binary condition differences to quantify the impact of a task feature (e.g., stimulus period vs. baseline period, high vs. low levels of conflict, familiarity vs. novelty, or houses vs. faces). The resulting implementation of GED (which we designate *comparison scanning GED*, csGED) identifies the frequencies at which two conditions are maximally separable, lending insight into differences in frequency content between conditions. Furthermore, csGED allows for exploration of the nature of the sources contributing to these frequency differences, by computing and analyzing source-specific time courses and topographies.

We aim to demonstrate the value of csGED to identify sources (i.e., spatial filters) of oscillatory frequencies that maximally differentiate two conditions. We quantified the accuracy of csGED source reconstructions in simulated data with a known ground truth, and we demonstrated the practical applicability of the method on a published MEG dataset. We thus show that csGED can reveal the frequency spectrum of condition differences in a data-driven way.

## Methods

### Description of problem statement and mathematical solution

M/EEG and LFP are subject to signal mixing effects. Such mixing complicates the relation of measured signals to cognition, as many functionally independent neural sources may contribute to the signal at any sensor. Because mixing is instantaneous and linear (Nunez & Srinivasan, 2006; Parra et al., 2005), multivariate linear source separation methods are able to approximate the constituent statistical sources, decomposing the mixed signal into linearly separable components. Blind source separation methods (PCA, ICA) are optimized to capture salient features in the full dataset, but this is not ideal for all purposes: in neuroscience, we are often interested in only a subset of the data, namely that related to a cognitive or behavioral phenomenon. We here validated a feature-guided source separation method based on generalized eigendecomposition (GED, also known as joint decorrelation; de Cheveigné & Parra, 2014). GED allows for the specification of features of interest in the data, which we here harnessed to tune the source separation to both condition differences and frequency content of the signal. To distinguish the general method from the specific condition-contrasting and frequency-scanning application in this work, we refer to our approach as *comparison scanning GED* (csGED).

The goal of csGED is to map the frequency spectrum of source differences between two conditions, designated A and B. The data in each condition must be broadband; it can be taken from subsets of trials in an experiment, or from time windows within trials. Data should be appropriately cleaned: noisy trials, subjects, sensors, and non-brain artifacts should be rejected or projected out, e.g., by ICA cleaning.

First, the sensor-level signal **X** (channels-by-time-by-trials) is filtered, through Morlet wavelet convolution (or any other suitable narrowband filter), at a range of frequencies *f* that span the frequency spectrum to be investigated. This filtering yields narrowband datasets **X**_f_, from which condition-specific narrowband subsets **X**_Af_, **X**_Bf_ can be extracted. To improve sensitivity, data can be restricted to time windows during which the condition differences are expected to be present; this should be done after filtering in order to avoid edge artifacts. In preparation for computing the covariance, the data in **X**_Af_, **X**_Bf_ are mean-centered by subtracting the average activity per trial from each time point in that trial, yielding a mean of 0 for each sensor.

Next, two sensor-by-sensor covariance matrices **K**_Af_ and **K**_Bf_ are constructed from the mean-centered narrowband data **X**_Af_, **X**_Bf_ for each frequency *f*. (eq. 1, see also Figure 1A):

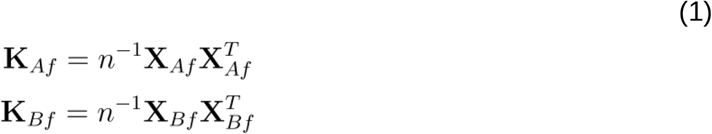

where *n* is the number of time points minus one in **X**_Af_ and **X**_Bf_, respectively, normalizing the covariance matrices with regard to duration (to facilitate comparison of eigenvalues for time windows of different length). Trials in **X**_Af_, **X**_Bf_ can be concatenated and used to compute the covariance matrix at once, or covariance matrices can be computed for each separate trial and averaged; we find that the latter option tends to produce cleaner covariance matrices with more stable solutions.

**Figure 1.**
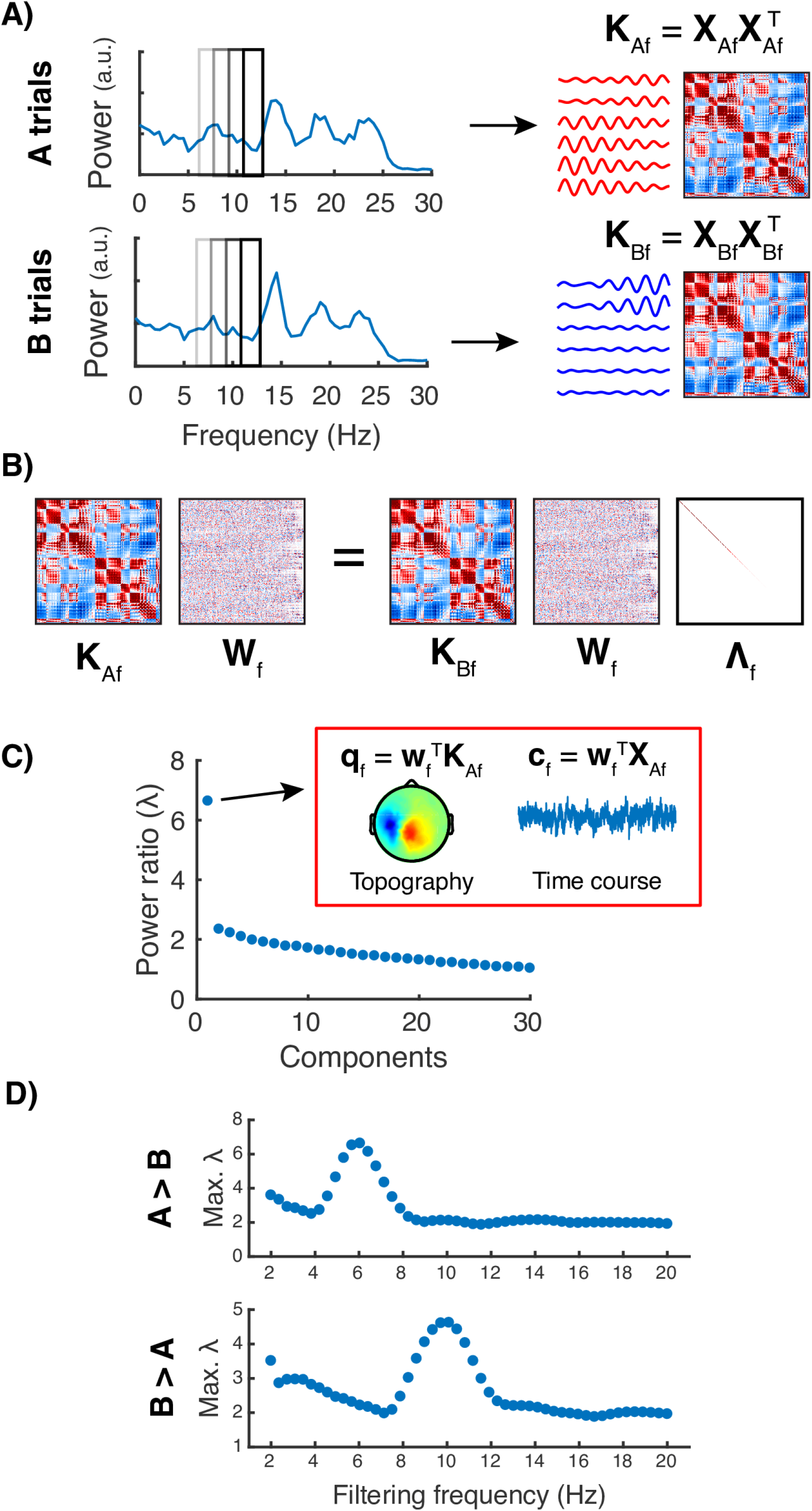
Illustration of the method as applied to simulated MEG data with unique signal and shared noise characteristics. **A)** Trial data from conditions A and B (**X**_A_ and **X**_B_) are filtered at specific frequencies f. Covariance matrices **K**_Af_, **K**_Bf_ are constructed from the narrowband data **X**_Af_, **X**_Bf_. **B)** Matrices **K**_Af_ and **K**_Bf_ are entered into the generalized eigenvalue equation, yielding a set of eigenvectors **W**_f_ and corresponding eigenvalues **Λ**_f_ that maximize the energy ratio between **K**_Af_ and **K**_Bf_ (or **K**_Bf_ and **K**_Af_, respectively). **C)** Each eigenvector (component) can be used to construct a component topography **q**_f_ and time course **c**_f_. The component with the largest eigenvalue λ is retained to construct a spectrum of maximum eigenvalues. **D)** The maximum eigenvalues at each filtering frequency f peak around 6 Hz when maximizing A > B and at 10 Hz when maximizing B > A, reflecting the ground truth signals present in conditions A and B respectively.

As GED finds sources that maximize the “multivariate power ratio” between two covariance matrices, one of the matrices must be chosen to be optimized (the *signal matrix*) and the other is chosen to be minimized (the *reference matrix*). To investigate two conditions of interest in csGED, the generalized eigenvalue equation is solved once using **K**_Af_ as signal matrix and **K**_Bf_ as reference matrix, and once with **K**_Bf_ as signal and **K**_Af_ as reference. The reference matrix can be shrinkage-regularized to improve separability and stability; details are provided in the next subsection.

By maximizing the Rayleigh quotient between **K**_Af_ and **K**_Bf_ (**K**_Af_ being signal and **K**_Bf_ being reference; eq. 2), a single set of weights **w**_Af>Bf_ is found. This **w**_Af>Bf_ captures the direction in which **K**_Af_ and **K**_Bf_ are maximally separable, i.e., where the power ratio between them is highest. The associated scalar value λ_Af>Bf_ (eq. 3) quantifies this power ratio. For our purposes, **w**_Af>Bf_ describes a spatial filter that captures the maximal energy ratio between conditions A and B filtered at frequency *f*. As such, it provides a spatial weighting that attempts to isolate the source that generates the most *f*-Hz energy in condition A compared to condition B. Conversely, maximizing the Rayleigh quotient between **K**_Bf_ and **K**_Af_ yields weights **w**_Bf>Af_ and power ratio λ_Bf>Af_, and isolates the source that generates most *f*-Hz energy in condition B compared to condition A.

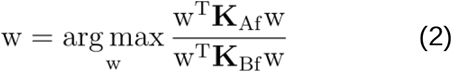

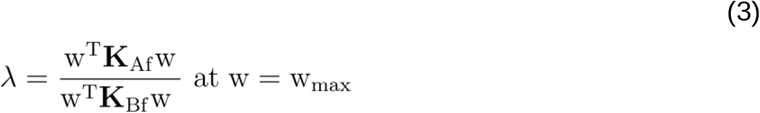

Equations 2 and 3 can be extended from a single vector computation to a matrix computation (eqs. 4 and 5; see also Figure 1B). The resulting generalized eigenvalue equations, when solved, each yield a matrix of weights **W** and a diagonal matrix of associated eigenvalues **Λ**.

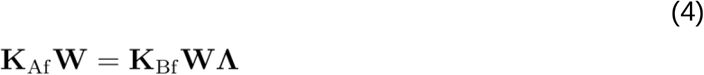

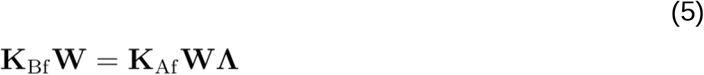

This generalized eigendecomposition equation is solved once for **K**_Af_ > **K**_Bf_ (eq. 4) and once for **K**_Bf_ > **K**_Af_ (eq. 5) at each frequency *f*. From each resulting **Λ**_Af>Bf_, **Λ**_Bf>Af_, the largest eigenvalue is extracted, as a measure of the maximum energy ratio between **K**_Af_, **K**_Bf_ (i.e., maximum separability between conditions) in that frequency. The associated set of weights (i.e., an eigenvector, or component) in **W**_Af>Bf_, **W**_Bf>Af_ is the spatial filter that attempts to isolate the source maximally separating the conditions at that frequency. To facilitate magnitude comparisons between components, extracted eigenvectors **w** are normed to unit length (eq. 6):

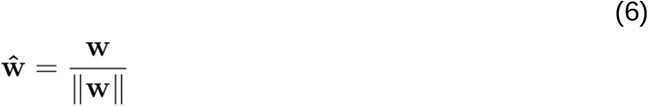

Plotting the largest eigenvalue found at each temporal filter frequency results in a frequency spectrum of separability, with peaks around frequencies where sources are found that strongly differentiate the conditions (Figure 1D). Components driving the spectral peaks can be further inspected by using the eigenvectors to compute component topographies and time courses.

#### Shrinkage regularization

The reference matrix can optionally be shrinkage-regularized (eq. 7) to improve separability and stability and, if necessary, correct potential rank-deficiency. Shrinkage regularization shifts the reference matrix towards a scaled version of the identity matrix (e.g., 1%), and thus shifts the GED procedure towards a PCA. In practice, 1% shrinkage improves separability for noisy or rank-deficient data, without affecting the decomposition of clean, full-rank data (Cohen, 2020). The signal matrix is best left intact to retain an accurate representation of the signal to be maximized.

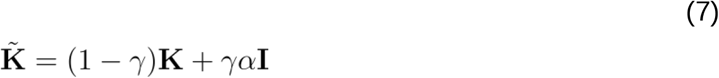

Here **K** is the original reference matrix, 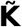 is the regularized reference matrix, γ is the degree of shrinkage (we used 0.01, which corresponds to 1%), **I** is the identity matrix of equal size to **K**, and α is the average of the eigenvalues of **K** (eq. 8):

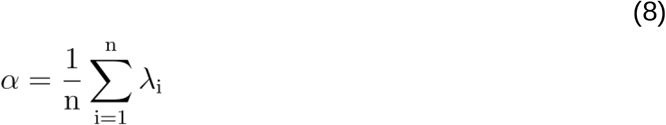

Following regularization, the regularized matrix is used in place of the original reference matrix in Equations 2 through 5.

#### Computing component topographies

From each component eigenvector **w**, a component topography **q** can be constructed as a linear combination of the covariance matrix weighted by the eigenvector (cf. (Haufe et al., 2014); see also Figure 1C):

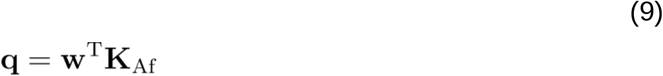

As such, the main difference between the eigenvector (spatial filter) and the scalp topography is that the scalp topography reflects not only the sensor weight, but also the patterns in the sensor covariance.

#### Computing component time courses

Component time series **c** can be constructed as linear combinations of the sensor-level time courses in **X**, weighted by the eigenvector **w** (see also Figure 1C):

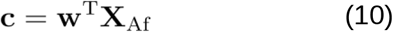

where **X** is the original (unfiltered, broadband) channels-by-time-by-trials data, and **c** is the component-by-time-by-trials result. Note that while **w** is computed from narrowband data, the fact that **X** is broadband results in the component time series **c** being broadband. The broadband nature of the component time series allows for time-frequency decomposition, which can serve both to verify that activity in **c** occupies a narrow range of frequencies, and to explore activity in other frequencies captured by the same spatial filter.

#### Time-frequency decomposition

Component time series can be further characterized through time-frequency decomposition. We implemented this by narrowband-filtering the time series **c** at a range of frequencies, through trial-by-trial convolution with complex-valued Morlet wavelets spaced throughout the frequency range of interest. Morlet wavelets were constructed as follows (eqs. 11 and 12):

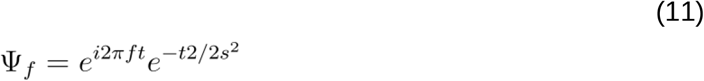

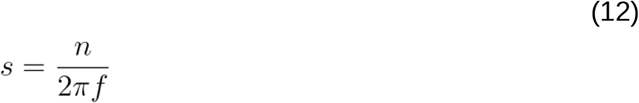

where *t* is time, *f* is frequency, *s* is the standard deviation of the Gaussian that modulates the complex sine wave, and *n* is the number of wavelet cycles. The number of cycles per wavelet governs the tradeoff between temporal and spectral precision. Both datasets considered in this paper were decomposed at 40 different frequencies. These frequencies were linearly spaced from 2 to 20 Hz for the simulated data and logarithmically spaced from 4 to 80 Hz for the empirical data. Wavelet widths were logarithmically spaced from 4 to 10 cycles.

Convolving the component time series with the created Morlet wavelets produces a (temporally smoothed) power time series for each frequency. To correct for the higher power observed at lower frequencies (Cohen, 2014), the signal power at each frequency is dB-scaled relative to the average baseline power at that frequency. Baseline windows should be relatively free from perturbations, last at least as long as one full cycle at the lowest decomposition frequency, and be far enough removed from trial epoch edges to avoid filtering-related edge artifacts. We selected a baseline window of 0.8 to 0.2 seconds prior to stimulus onset for the simulated data, and 0.8 to 0.2 seconds prior to fixation cross onset for the empirical data.

### Simulations

Before analyzing empirical data (where the ground truth is unknown), we validated and explored our methods in simulated data, in which the ground truth is known and where it is possible to arbitrarily control the signal-to-noise ratio (SNR). To this end, we simulated MEG data for two conditions (“A” and “B”) with unique signal and shared noise characteristics. The csGED procedure was applied once for A > B and once for B > A. Peaks in the eigenvalue spectra were identified, and the peak-associated component time series were compared to the ground truth signals (by computing shared spatial variance R2), yielding a measure of reconstruction accuracy. This reconstruction accuracy was compared to the accuracy at best sensor for each dipole, i.e., the sensor to which the dipole projected most strongly (positively). We varied the signal amplitude relative to the noise amplitude to evaluate results for different SNR.

#### Dipole selection

A model containing 15000 cortical surface-level dipole locations, and each dipole’s associated projection weight matrix (leadfield matrix) to sensor locations corresponding to the CTF 275 system, was generated using Brainstorm (Tadel et al., 2011). As a detailed anatomical representation of the cortex was not necessary, the model was downsampled to 3000 dipoles, improving simulation speeds. The leadfield projection weights for each dipole were normalized by that dipole’s maximum projection weight across sensors, so that all dipoles projected to the scalp comparably strongly (eq. 13):

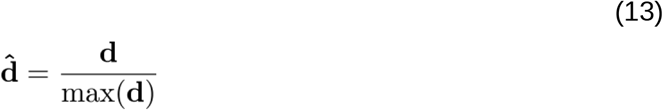

where **d** are the original and 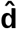 are the normalized projection weights for a single dipole.

To demonstrate the efficacy of csGED in separating sources with partially overlapping topographies, we randomly selected two signal-generating dipoles such that their topographies had a spatial correlation of R2 = 0.4. One of those dipoles was active in condition A trials and the other was active in condition B trials. Dipole projection topographies are shown in Figure 2A.

**Figure 2.**
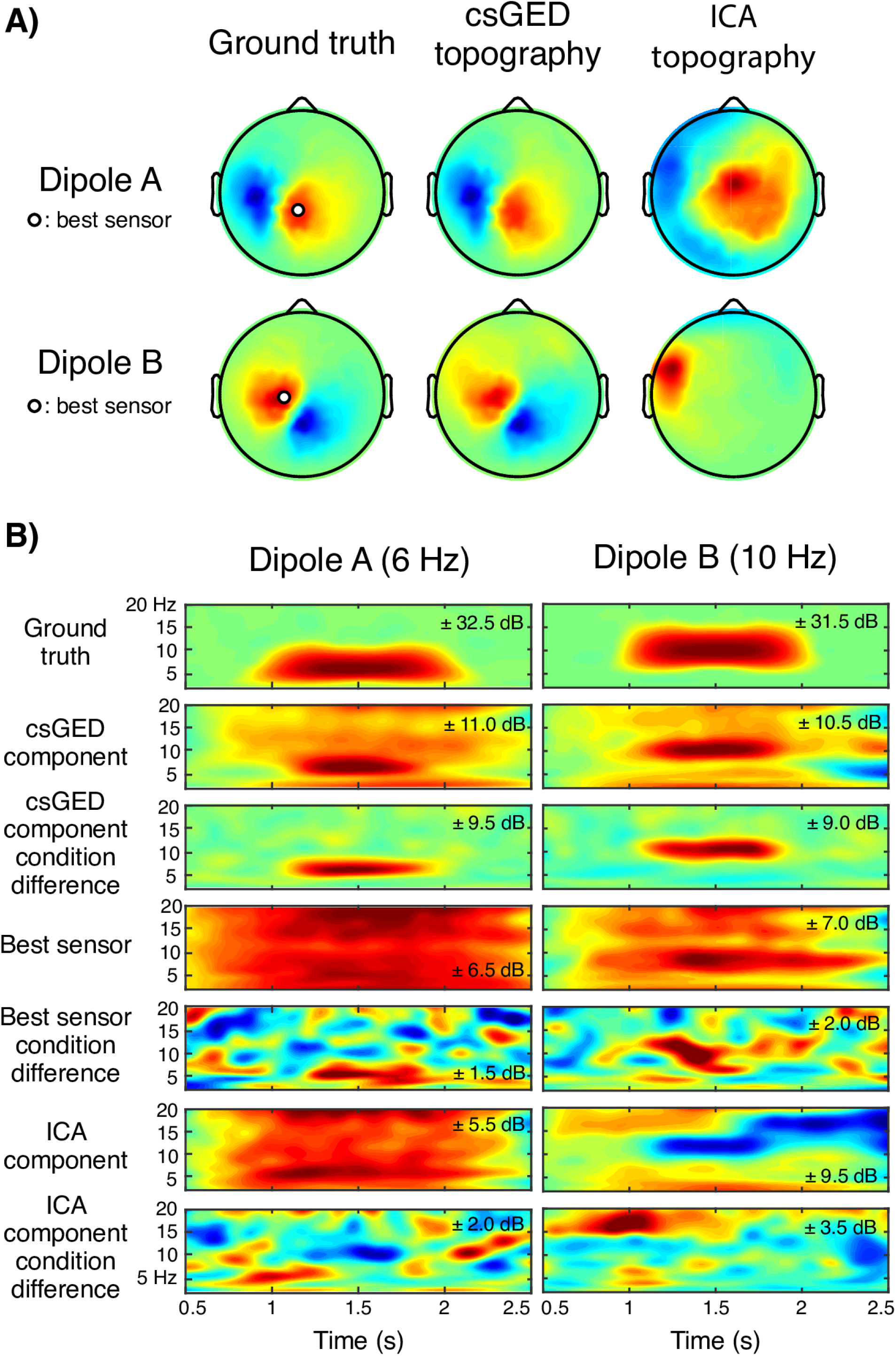
Ground truth signals and signal reconstructions in simulated MEG data. **A)** Left: ground truth scalp projections of the signal-generating dipoles in both conditions. Middle: GED-reconstructed topographies of the signal dipoles for SNR 10^−5^ are highly similar to the ground truth topographies (shared spatial variances R^2^ = 0.97 and 0.94 for dipole A and B respectively), outperforming ICA-reconstructed topographies (right; shared spatial variances R^2^ = 0.43, 0.22). **B)** Time-frequency (TF) decompositions illustrate that the GED component time series better approximate the ground truth signals (temporal R^2^ = .031, .049 for dipole A, B) than the signals at the best sensors (temporal R^2^ = .006, .008) or the ICA reconstructions (temporal R^2^ = .001, .000). Condition differences (third, fifth, seventh rows) were obtained by subtracting TF-decomposed A trials from B trials and vice versa. The GED component condition difference provides a particularly close approximation of the ground truth signal.

In addition to these signal dipoles, a random set of 1000 “distractor” dipoles (spread throughout the cortical model) was selected for each simulation, of which a randomized 30% on average would be active on any given trial. These distractor dipoles simulated noise (i.e., task-irrelevant activity) within the brain. A signal dipole could not also be a distractor dipole.

#### Signal and noise generation

We simulated 200 3-second trials, of which the first 100 represented condition A and the last 100 represented condition B. Stimulus presentations were simulated during the 1– 2 s time window. This meant that on each trial in condition A, the first signal dipole was active during this time window while the second remained quiet. This was reversed for trials in condition B, where the second dipole was active while the first remained quiet. The activity each dipole generated was the time-domain representation of a uniformly randomly generated frequency spectrum, bandpass-filtered (through frequency-domain Gaussian convolution, Gaussian FWHM of 2) at a given frequency. Signal frequencies were chosen to be 6 Hz for dipole A and 10 Hz for dipole B.

On each trial, each of the 1000 distractor dipoles had a 30% chance of being active. An active dipole generated a waveform centering on a randomly chosen frequency with a maximum of 25 Hz, during a randomized time window of >0.3 s duration. Time windows were re-randomized on each trial, ensuring that any spurious phase coherence between dipoles did not persist across trials. A tapered cosine window with a fraction of .6 was applied to the dipole waveforms so that they faded in and out, preventing sharp edges from causing filter artifacts. The waveforms were scaled to a fixed root mean square (RMS) amplitude of 3. In addition, white noise with a fixed SD of 0.5 was added to all (signal, distractor, and inactive) dipoles for the entire duration of the trial. The total activity for each dipole was projected to the scalp through multiplication with the leadfield matrix, resulting in a mixed sampling of dipole activity at each of the 275 simulated sensors.

To vary the SNR, the simulated signal was scaled to a specific RMS amplitude. The SNR for a given simulation was computed as the power ratio between total signal dipole amplitude and total distractor dipole amplitude (eq. 14), accounting for the fact that distractor dipoles were on average 300 times as plentiful as signal dipoles on a given trial and that scalp projection magnitudes for all dipoles were normalized. White noise amplitudes were not taken into consideration for the SNR value. Note that the SNR reflected only “brain signal” and “brain noise”, as we did not attempt to model other sources of noise.

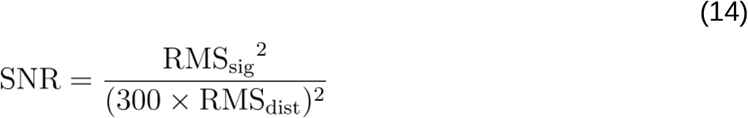

Here RMS_sig_, RMS_dist_ are the root mean square amplitudes for the signal and distractor dipoles, respectively. Note that signal and noise dipoles were distributed throughout the brain. Thus, the SNR at the topographical level reflects a mixture of all dipoles and thus in practice varies between sensors.

#### Contrasting conditions at each frequency

The previously described GED procedure was applied to the simulated data. The scalp-level signal **X** (channels-by-time-by-trials) was filtered at each frequency *f* using Morlet wavelet convolution, at 50 frequencies linearly spaced from 4 to 20 Hz. A time window of interest (1–2 s) was extracted from each trial, exactly matching the simulated stimulus duration. Covariance matrices **K**_Af_ and **K**_Bf_ were constructed from the A and B trials, and the eigendecomposition equation was solved. From the largest eigenvalues, a frequency spectrum was constructed, allowing us to derive at which frequencies the power ratios for A > B and B > A were maximal (Figure 1D).

#### Selecting frequencies of interest

To identify the components of interest, we first identified the frequencies of interest. We selected eigenvalue peaks in the frequency spectrum to determine at which frequencies A > B and B > A were maximal. An eigenvalue in the spectrum that was higher than the eigenvalues at both neighboring frequencies was considered a peak, with no additional constraints. The single highest peak in each spectrum was selected, and the corresponding eigenvector was dubbed “component A” for the A > B spectrum, and “component B” for the B > A spectrum. These components were used to construct component time series, which were then time-frequency decomposed for visual inspection, as described earlier.

#### Comparing reconstruction accuracy

We compared the GED-derived reconstruction of the ground truth dipole signal to two other types of reconstruction: the signal at one sensor, and an ICA component. For the single-sensor signal, the simulated MEG sensor to which each signal dipole projected most positively was selected as the most representative (“best”) sensor for that dipole. The ICA reconstruction was created by applying the JADE algorithm (constrained to 30 ICs) (Cardoso & Souloumiac, 1993) to the broadband data from each condition, and selecting the top component.

Reconstruction accuracy was quantified as the shared temporal variance (R^2^, squared correlation coefficient) between the ground truth dipole time series and the reconstructed time series in the relevant condition. Simulations were repeated five times with the same signals and different noise, and results were averaged over iterations and over A > B and B > A contrasts. R^2^ values were plotted against the simulation SNR, to compare the accuracy of the reconstruction methods in the presence of noise.

### Empirical data

We selected a previously published dataset (Dijkstra et al., 2018) to analyze. Grounds for selection included the type of data collected (MEG, suited to source separation) and the binary task condition difference (perception vs. imagery of visual stimuli). Furthermore, imagery data can be challenging to analyze, as it is weakly time-locked to external stimuli or instructions (Dijkstra et al., 2019; Vidaurre et al., 2019). The absence of time-locking is not an issue for GED, as the method harnesses spatial covariance rather than temporal characteristics (beyond the selection of a general time window to analyze) to identify spatial filters. It is thus well-suited to the analysis of fluctuating neural dynamics, such as those exhibited during imagery.

#### Subjects

30 human subjects with normal or corrected-to-normal vision participated in this study. Subjects provided written informed consent. The Committee Human Research (CMO) Arnhem-Nijmegen approved the study, and all research was conducted according to the appropriate guidelines. Five subjects were rejected (for movement, failure to follow task instructions, or technical problems), leaving 25 subjects.

#### Task design

Full details are available in the original publication (Dijkstra et al., 2018); we here provide a brief summary that is relevant for our analyses. Subjects were sequentially shown two stimuli (face and house, or house and face; 0.8 s each, taken from a set of 8 possible stimuli), and were then cued to imagine (3.5 s) either the first or second stimulus shown to them. Visual stimuli were separated by a random 0.4–0.6 s interval. The visual stimuli were separated from the imagery cue by a 0.5 s noise mask and another 0.4–0.6 s interval. Subjects completed 240 trials (120 per category; stimulus order counterbalanced) in blocks of 10.

#### Data acquisition

MEG data were acquired in a shielded room using a whole-head 275 axial gradiometer MEG system (MEG International Services Ltd., Coquitlam, BC, Canada) at a sampling rate of 1200 Hz. Five sensors were defective and recorded no data. Head location was monitored and corrected during between-block breaks. Electro-oculograms (EOG), an electrocardiogram (ECG), eye position, and pupil size were recorded to facilitate the removal of eye- and heart-related artifacts.

#### Data cleaning and preprocessing

Data were preprocessed using MATLAB (MATLAB 2014a, The MathWorks, Natick, 2014) and the FieldTrip toolbox (Oostenveld et al., 2011). Data were high-pass filtered at 0.5 Hz. 50 Hz line noise, along with bandstop-filtering-resistant artifacts at 50 Hz, was successfully filtered out using the ZapLine method (de Cheveigné, 2019) by removing a single noise component from each trial. Trial epochs were defined as lasting from 1 s prior to the appearance of the fixation cross (2 s prior to stimulus onset) to 1 s after the imagery time window, for a total of 15 s. Trial rejection was based on Dijkstra et al., 2018: if any of the relevant time windows (first visual stimulus, second visual stimulus, or imagery) had been rejected in the original study, we here rejected the full trial. After trial rejection, 192.4 ± 34.3 (mean ± SD) trials remained for each of the 25 subjects (minimum of 98 trials), each of which included a perception (P) and an imagery (IM) time window.

After trial rejection, data were downsampled to 300 Hz and independent component analysis (ICA) was performed. ICs capturing clear heart and eye artifacts were removed, based in part on correlations with the ECG/EOG/eye-tracking data and in part on visual inspection of the IC topographies. Data were scaled from Tesla to picoTesla prior to analysis, to prevent precision loss from rounding errors.

#### Contrasting conditions at each frequency

The csGED procedure was performed separately on each subject. First, the full-length trials were filtered at a range of frequencies through Morlet wavelet convolution. To optimize the trade-off between frequency resolution and computation speed, we first performed low-resolution scans with a frequency interval of 2 Hz (from 2 to 80 Hz), and then re-scanned in an 8 Hz range (4 Hz on either side) around identified peaks, with a frequency interval of 0.5 Hz.

Next, at each frequency, three relevant task windows were extracted from the full-length trial: one during the presentation of the first visual stimulus (“perception 1”, P1; 0.8 s duration), one during the second visual stimulus (“perception 2”, P2; 0.8 s duration), and one during the imagery window (IM; 3.5 s duration). To allow the GED to optimize for general perception-related processes, P1 and P2 were concatenated into a single “perception” (P) window. Sensor-by-sensor covariance matrices **K**_Pf_, **K**_IMf_ were computed per trial and then averaged. Then, for each subject, the eigendecomposition equation was solved once to find spatial filters that maximized **K**_Pf_ > **K**_IMf_, and once for **K**_IMf_ > **K**_Pf_. Each solution yielded a set of eigenvalues and eigenvectors, of which the largest eigenvalue and corresponding eigenvector were retained and the others were discarded.

#### Selecting frequencies of interest

Subject-specific eigenvalue spectra typically contained multiple peaks. Local maxima in each spectrum were selected by identifying eigenvalues that 1) were flanked by lower eigenvalues at neighboring filter frequencies, 2) had a prominence (a measure of peak independence, defined as a peak’s height relative to the lowest neighboring trough, “neighboring” meaning there is no higher peak separating the two) of at least 0.01 λ (chosen based on visual inspection), and 3) were separated by at least 2 Hz (chosen based on visual inspection). If multiple eigenvalue peaks fell within 2 Hz of each other, the largest eigenvalue was selected. Selected eigenvalue peaks indicated a locally maximal power ratio for P > IM or IM > P at that frequency, meaning a spatial filter specific to that frequency was found that strongly maximized energy in one condition and minimized energy in the other. Components at the peaks were used to compute component topographies and component time series as described previously. Component time series were time-frequency decomposed for visual inspection, as described earlier.

#### Group-level analysis

At this point, two condition-specific sets of eigenvalue peaks had been identified for each subject, and the peak-associated components could be inspected in terms of topography and time-frequency characteristics. This concludes the core csGED procedure. However, as we observed nonnegligible variability in peak frequencies between subjects (Figure 4A), we additionally analyzed the components at the group level to facilitate the interpretation of divergent results.

The next subsections address the pooling of components across subjects. We here operated on the assumption that components that 1) occupied the highest eigenvalue (i.e., were most salient) and 2) were close in frequency would be functionally similar between subjects, and could thus be averaged. This assumption reflects common practice in electrophysiology, where multi-subject time-frequency measurements are typically averaged without regard for linear separability. However, other criteria could be used to group components together, and users of the csGED method are encouraged to consider the best criteria for their use case.

#### Kernel density smoothing

To reconcile variability in the subject eigenvalue spectra and the extracted peak frequencies, we computed a kernel density estimate from the previously selected peaks per individual subject. Kernel density estimates are useful because they remove the biases resulting from different raw eigenvalue magnitudes across subjects and across frequencies. By applying this method of combining components, we made the assumption that components in different subjects are functionally similar when 1) they occupy the largest eigenvalue and 2) the frequencies to which they most contribute are close together.

Kernel density estimation was performed by taking the eigenvalue peak frequencies from all subjects, representing each as a Gaussian with a height of 1 and SD of 2 Hz (chosen to create a reasonably smooth plot), and summing over Gaussians. This procedure is illustrated in Supplementary Figure S1. The resulting kernel density functions (KDFs) reflected a combination of the proportion of individual-subject peak components around a given frequency, and how close to that frequency the peak components were located.

#### Averaging over components

Upon visual inspection, the KDFs exhibited multiple salient peaks (subjectively defined as exceeding 0.01, a.u.), reflecting frequencies near which many subjects presented with eigenvalue peaks. To facilitate comparing component characteristics, we averaged together component time courses, topographies, and time-frequency decompositions based on the components’ distance to the KDF peak frequency. Specifically, frequency boundaries around each KDF peak were defined so that the ratio between the center frequency and the frequency band width was 2.33 (following the “golden rule”; Klimesch, 2018). Each subject’s peak component closest to the KDF center frequency was included in the average, as long as it remained within frequency boundaries. As such, each subject contributed a maximum of one component; a given peak average might therefore not include components from all 25 subjects. In practice, 16.8 subjects on average contributed components to a KDF peak, with a minimum of 7 and a maximum of 23.

### Code and data accessibility

The custom-written MATLAB analysis code has been made available on GitHub at http://www.github.com/marrit-git/csGED. The empirical data are available from https://data.donders.ru.nl/.

## Results

### Simulation results

At a SNR of 10^−5^, equivalent to placing a single signal dipole among 300 distractor dipoles generating sine waves of equal amplitude, the csGED-derived eigenvalue spectra exhibited clear peaks centering on the frequency of the active dipole in that condition (Figure 1D). The csGED procedure thus successfully identified the frequencies at which covariance power in condition A was maximal compared to condition B and vice versa, without prior expectations about these frequencies.

The components corresponding to the eigenvalues at these maxima represented the GED-derived reconstruction of the original dipole signal as a linear mixture of sensor data. At a SNR of 10^−5^, the topographical reconstructions (spatial filters) were highly similar to the original dipole projections (Figure 2A; squared spatial correlation R^2^ = 0.97 and 0.94, respectively), surpassing ICA-derived topographical reconstructions (R^2^ = 0.43 and 0.22; *p* = 0.014, one-sided *t*(2) = 5.93).

Time-frequency decompositions (Figure 2B) allowed for visual comparison of the original time courses to the reconstructions. At a SNR of 10^−5^, the reconstructed time courses provided a closer fit to the ground truth signal (temporal R^2^ = .031, .049 for dipole A and B, respectively) than the signal at the best sensor (though not significant; temporal R^2^ = .006, .008 for A and B; *p* = 0.033, one-sided *t*(2) = 3.64, Bonferroni-corrected α = 0.025) or the ICA time series (temporal R^2^ = .001, .000 for A and B; *p* = 0.024, one-sided *t*(2) = 4.38, Bonferroni-corrected α = 0.025).

The low R^2^ values of the GED components are attributable to the noise and distractors in other frequencies: narrowband-filtering all signals at 6 and 10 Hz (FWHM 2), respectively, provided closer fits (R^2^ = .502 and .417) for the GED-derived time courses, though still only R^2^ = .022 and .071 for the sensor-level signals, and R^2^ = .000, and .000 for the ICA components. Taken together, these results show that at this specific SNR, source separation components captured the ground truth brain signal better than single-sensor analyses or ICA did.

Next, we quantified the influence of the SNR on the reconstruction accuracy of all three methods. The SNR was parametrically varied by changing the amplitude of the signal dipole. The GED-derived reconstructions outperformed both the best sensor and ICA at reconstructing a given signal, for all but the lowest and highest SNRs (Figure 3).

**Figure 3.**
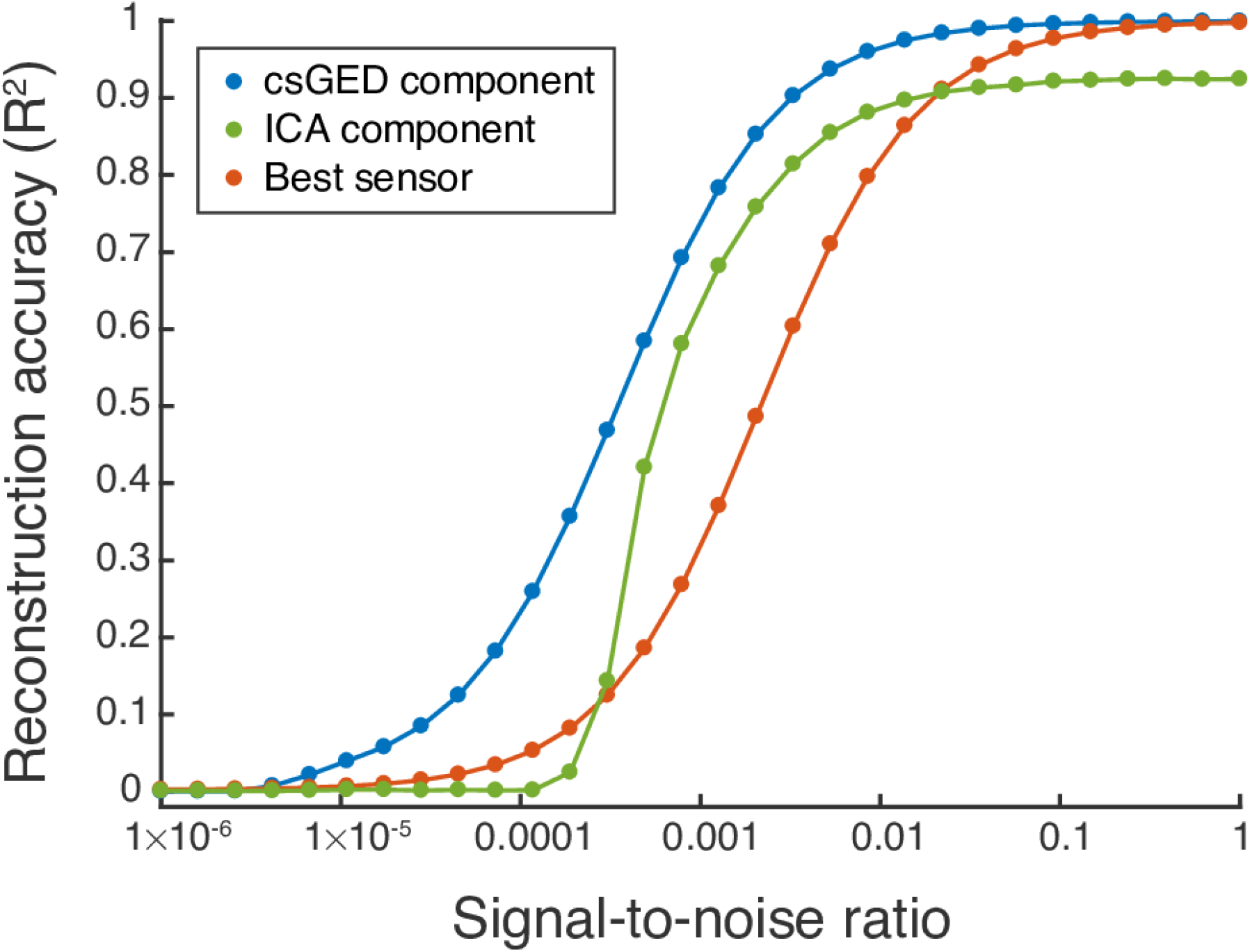
In simulated MEG data, the csGED-derived component time series captured the ground truth more accurately than either the best sensors or the ICA-derived component time series, even in the presence of large amounts of noise (300 noise dipoles for 1 signal dipole).

Together, these simulation results demonstrate that csGED successfully identified the frequency content that maximally differentiated task conditions, without prior expectations about the relevant frequencies. Furthermore, the ground truth signal at a given frequency was approximated more accurately with this multivariate analysis than with two common alternatives: single-sensor analysis and ICA.

### Empirical results

Each trial in our empirical dataset contained a time window during which subjects perceived visual stimuli (P) and a time window during which subjects imagined one of the stimuli (IM). GEDs contrasting the P and IM time windows on each trial were applied to data filtered at a range of frequencies, effectively scanning the frequency spectrum for high eigenvalues. This scan was performed once at low resolution (from 2 to 80 Hz with frequency intervals of 2 Hz) and then repeated at high resolution (frequency intervals of 0.5 Hz) around identified peaks, as described in the Methods section. The csGED method successfully identified eigenvalue peaks in the P > IM and IM > P frequency spectra for each subject, reflecting linearly separable components that maximally separated the P and IM conditions at those frequencies. P > IM and IM > P frequency spectra for individual subjects were variable (Figure 4A), likely reflecting intra-individual variability in peak oscillatory frequency.

**Figure 4.**
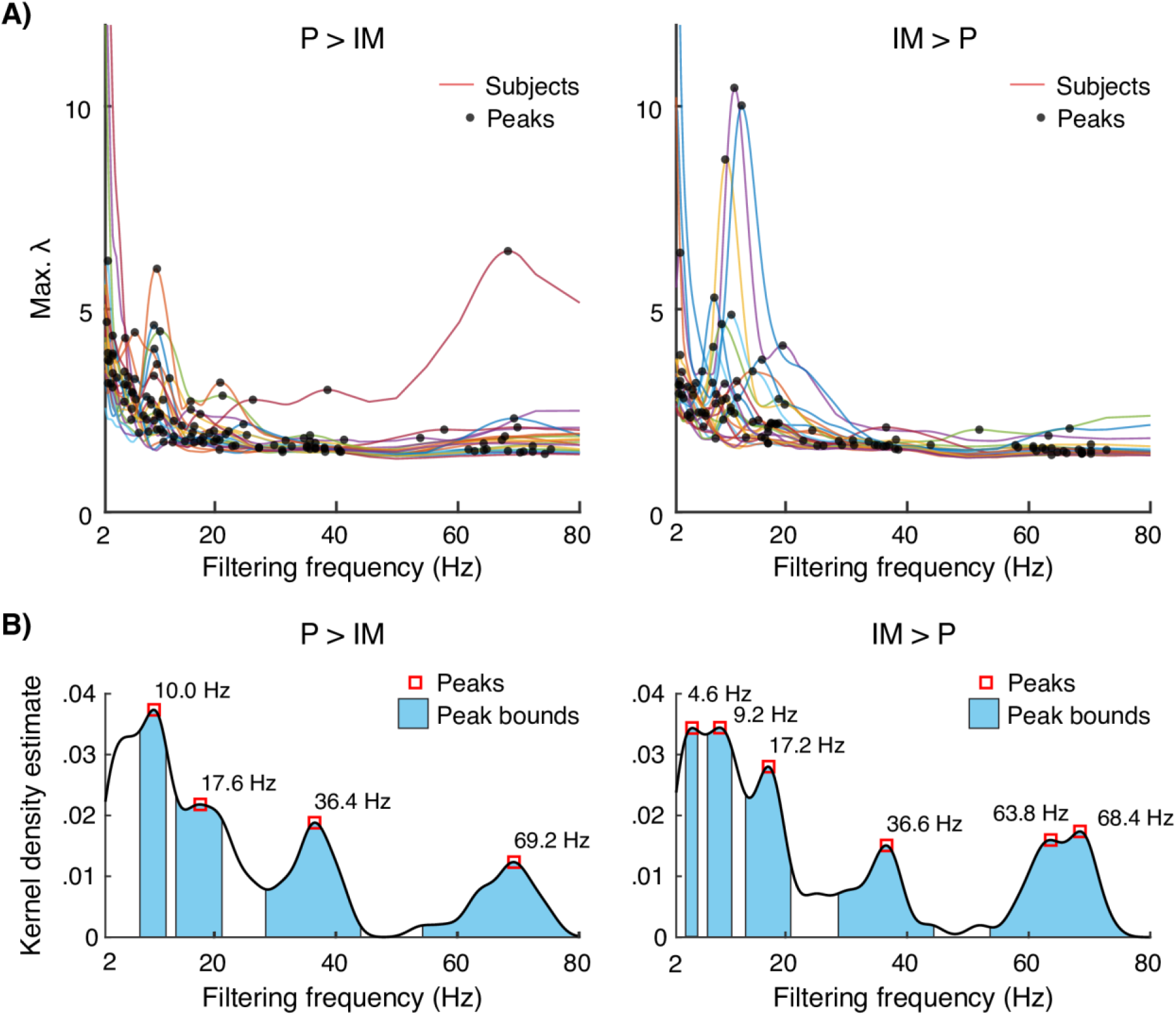
Results of condition-specific spectral scanning on empirical MEG data, contrasting perception (P) against imagery (IM) time windows. **A)** Eigenvalue spectra for all subjects. Peaks were identified and transformed using a KDF. **B)** Kernel density estimates show multiple peaks, with each peak indicating that multiple subjects had high-eigenvalue components around that frequency. Notably, peaks for P > IM and IM > P were observed at similar frequencies, indicating condition-differentiating, spatially dissociable networks, resonating at nearly identical frequencies. Blue marks the frequency boundaries for each peak, which were later used to constrain averaging across components. IM > P gamma frequency ranges (with peaks at 63.8 and 68.4 Hz) overlapped substantially (sharing 22 out of 23 components) and were merged together.

In order to combine data across subjects, a KDF was constructed from all eigenvalue peak frequencies, as illustrated in Supplementary Figure S1. Local maxima in the KDF (Figure 4B) indicate the frequencies at which subjects commonly exhibited an eigenvalue peak, while ignoring peak magnitude. KDF peak frequencies are listed in Table 1. The spectra exhibited peaks at similar frequencies for both condition contrasts. As the GED explicitly normalizes out commonalities between the conditions, this observation suggests that sources with different spatiotemporal characteristics, i.e., different topographies and activity patterns, were active at the same frequencies in the two conditions.

**Table 1.** Overview of peak frequencies extracted from the KDF, and summary statistics for the components selected as contributing to each peak frequency.

| Comparison | KDF peak frequency | Component frequency<br>(mean ± SD) | Component $n$ |
| --- | --- | --- | --- |
| P > IM | 10.0 Hz | 10.2 ± 0.95 Hz | 21 |
| | 17.6 Hz | 17.8 ± 2.09 Hz | 18 |
| | 36.4 Hz | 36.0 ± 3.07 Hz | 20 |
| | 69.2 Hz | 68.4 ± 3.96 Hz | 17 |
| IM > P | 4.6 Hz | 4.4 ± 0.49 Hz | 7 |
| | 9.2 Hz | 9.5 ± 1.09 Hz | 15 |
| | 17.2 Hz | 17.4 ± 1.43 Hz | 17 |
| | 36.6 Hz | 34.9 ± 3.30 Hz | 13 |
| | 66.6 Hz | 64.5 ± 4.70 Hz | 23 |

KDF-peak-related component topographies and time-frequency decompositions were averaged together for visual inspection. To do so, we selected components presumed to contribute to each KDF peak, by identifying the component for each subject that was closest to the KDF peak as well as within frequency band boundaries (specified as 2.33 x peak frequency, following Klimesch, 2018). Table 1 lists the number of components included per peak and the average component frequencies. Individual components included in select peak averages (alpha, 10.0 and 9.2 Hz; and gamma, 69.2 and 66.6 Hz) are shown in Supplementary Figures S2 through S5. Note that we here operated on the assumption that components occupying the highest eigenvalue and centered around the same frequencies are functionally similar, and thus can be meaningfully averaged.

Individual components, as well as component averages for each peak, varied both in topography and in time-frequency characteristics (Figure 5; Supplementary Figures S2 through S5), suggesting that csGED captured functionally diverse networks. Notably, csGED was able to harness condition differences to separate sources with overlapping spatial projections and frequencies (e.g., the P > IM and IM > P alpha components for subjects 1, 7, 15, and 19 in Figures S2 and S3). Conversely, it was able to harness frequency differences to separate spatially overlapping sources active in the same condition (e.g., the P > IM components for subjects 4 and 9 in alpha and gamma; Figures S2 and S4). Note that individual gamma components may contain alpha activity whereas alpha components may not contain gamma activity, and vice versa. This is because different sets of sensors are involved in the construction of the alpha and gamma component.

**Figure 5.**
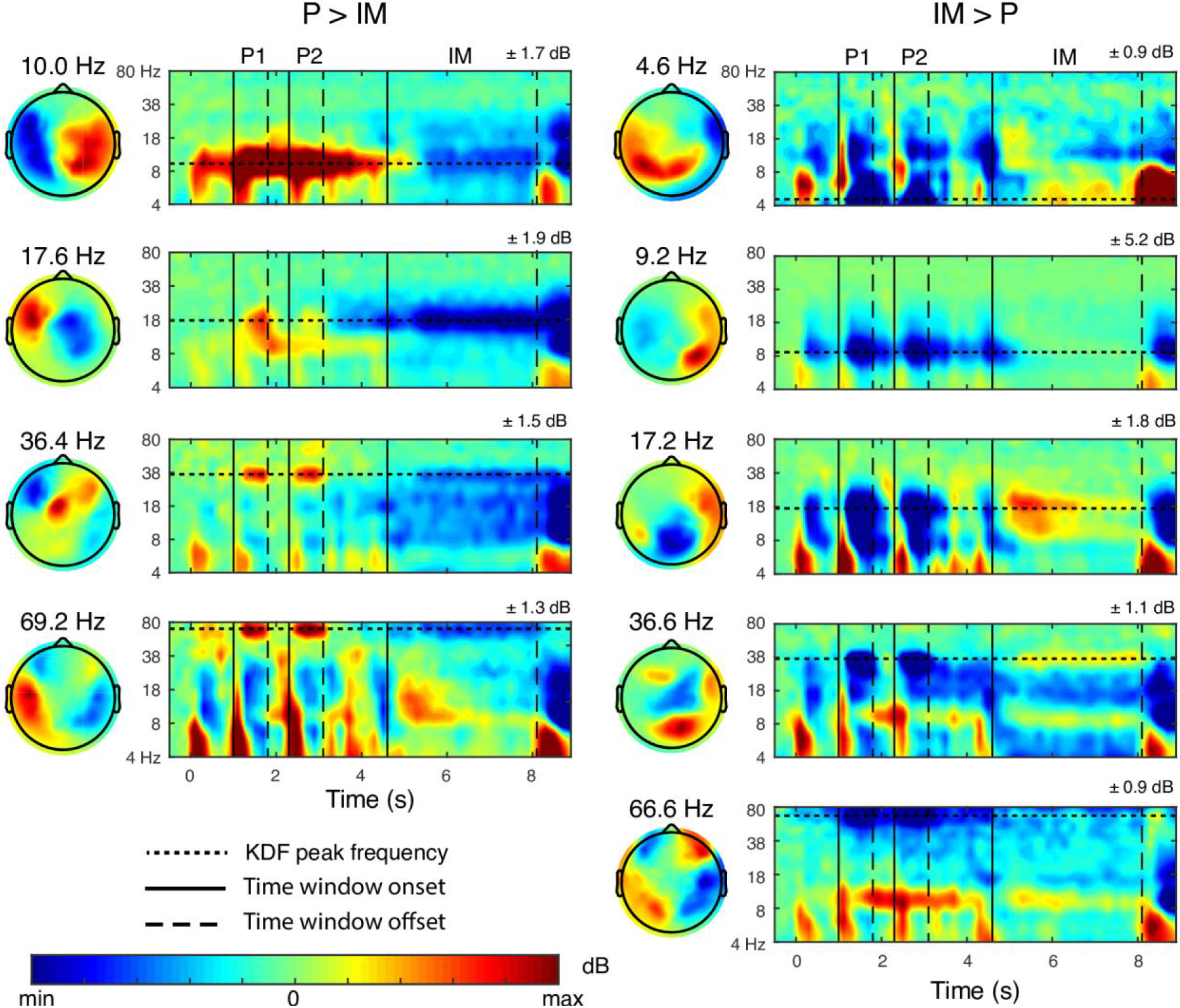
Average component topographies and time-frequency decompositions, encompassing each subject’s peak component closest to the overall KDF peak within frequency bounds. Average time-frequency decompositions tended to show power increases during the relevant condition and decreases during the other condition, indicating that csGED optimized for both.

Averaged component time-frequency dynamics (Figure 5) tended to show power increases during the condition of interest as well as power decreases during the other condition. This appeared to be a result of averaging, as individual component dynamics (Figures S2 through S5) tended to exhibit either power increases or power decreases, and rarely showed both. Many of the observed source dynamics were most pronounced during the perception time windows. Several features of these components are commonly linked to perception, such as short-lived alpha power decreases during perception (e.g., Fries et al., 2008; Klimesch et al., 2007) and gamma power increases during perception (Fries et al., 2008; Hoogenboom et al., 2006; Sedley & Cunningham, 2013). However, the persistent increases in alpha power and the decreases in gamma power are not typically observed in perception tasks. These may reflect perception-related source dynamics that are normally obscured, or they may reflect other processes; for example, the alpha increases may be related to attentional orienting (Thut et al., 2006). Gamma power decreases are rarely reported, regardless of task, and may typically be obscured by concurrent gamma power increases.

Dynamics during the imagery time window generally remained closer to baseline than dynamics during the perception time window. The identified IM > P sources were thus characterized more by suppression during perception than by imagery-related power changes. It is possible that csGED recovered sources driven by imagery-related dynamics, but that these spatial filters were associated with smaller eigenvalues and were therefore not included in our analyses.

## Discussion

### Summary

We here presented csGED, an analysis method that scans the frequency spectrum for sources that maximally differentiate two task conditions. While GED as a general method has been validated for different applications and on different types of neural data (Cohen, 2017a; de Cheveigné & Arzounian, 2015; de Cheveigné & Parra, 2014; Nikulin et al., 2011), this study is the first to concurrently combine frequency- and condition-sensitivity.

We first validated the source separation performance of csGED in simulated MEG data, for which the ground truth is known. We specified the operating frequency of signal-generating dipoles that were active in different conditions. csGED correctly identified the frequencies that differentiated the conditions, without prior hypotheses about these frequencies. Next, we examined the top sources that csGED found at these frequencies and quantified how accurately the sources captured the ground-truth signals. This source reconstruction accuracy was compared to the accuracy at the best single sensor for each signal (which is commonly taken to be representative for phenomena detectable in M/EEG; e.g., Asanowicz et al., 2019; Tiedt et al., 2016). For all but the highest and lowest signal-to-noise ratios, the csGED source reconstruction was markedly more accurate than the single-sensor representation of the source.

We next demonstrated the practical applicability of csGED in an empirical MEG dataset that compared visual perception and mental imagery (Dijkstra et al., 2018). As a result of its combined condition- and frequency-specificity, csGED was able to separate sources with overlapping topographies even when these sources generated activity at the same frequency or in the same condition. In sensor-level analyses, these separable sources would have remained linearly mixed (and thus confused) on account of their spatial co-occurrence. Furthermore, time-frequency decomposition of the source time series confirmed the presence of established neural signatures of perception in the source dynamics, suggesting that linear separation resulted in functionally meaningful segmentation.

In sum, csGED is able to successfully index the frequency spectrum of condition differences, and can accurately reconstruct signal sources whose dynamics may then be further inspected. We next highlight several features, idiosyncrasies, and limitations that may be relevant to future applications of csGED.

### Starting from clean data

csGED-identified components are not guaranteed to isolate brain sources. To prevent components from being contaminated by artifacts, data should be appropriately inspected and cleaned before applying the csGED method. Particular care should be taken to remove artifacts that affect sensor covariance differently in the data used for the signal matrix than in the data for the reference matrix, such as a noise dipole that is present in one condition and absent in the other (e.g., if there are more eyeblinks in one condition than in another). On the other hand, artifacts that equally affect the signal and reference matrix, such as line noise, will cancel out.

### The interpretation of sources

It should be kept in mind that csGED is performed without any anatomical constraints or biases; thus, isolated statistical sources may, but do not necessarily, correspond to individual, spatially restricted, anatomical generators. As csGED is a covariance-based method, the spatial filters found may reflect distributed networks that generate covarying signal. Furthermore, the *inverse problem* applies: multiple combinations of dipoles exist that can drive a given spatial distribution. For these reasons, we prefer more conservative statistical and topographical interpretations over anatomical speculations.

One assumption underlying csGED is that the sources are covariance stationary, that is, that the covariance structure is constant over the selected time windows and over repeated trials. Violations of this assumption—for example through head movements or traveling waves—may decrease reconstruction accuracy or may be captured as two statistical sources. We have previously investigated this possibility in empirical data (Zuure et al., 2020) and found it unlikely to be a significant issue.

### Other spatial filter methods

There are several spatial filtering methods used in electrophysiology research, including principal components analysis (PCA) and independent components analysis (ICA). Owing to different assumptions, different methods highlight different features of a dataset. csGED differs from unsupervised methods like PCA and ICA primarily in that it is hypothesis-driven, i.e., optimizes for a given condition contrast at a set range of frequencies. This optimization essentially maximizes the signal-to-noise ratio for that contrast (de Cheveigné & Parra, 2014) and lends GED increased distinguishing power for hypothesis-relevant sources. In an entirely hypothesis-free exploration, PCA and ICA may be better suited to detect salient sources of activity. However, it is rarely the case that even exploratory analyses cannot be targeted towards a contrast, such as a task vs. baseline window. We have previously found that GED reconstructs simulated sources with higher accuracy than PCA or ICA (Cohen, 2017b), lending further validity to the csGED method.

### Overfitting and underfitting

csGED is performed through the repeated solving of the generalized eigenvalue equation: once at every frequency. In this work, we have chosen to retain only the source with the largest eigenvalue (corresponding to the spatial filter explaining the most variance) from each of these solutions. The spectrum of separability was then constructed from these largest sources. Selecting a single source at each frequency and contrast carries risks of overfitting and underfitting: it is possible that we retain a source where in reality there is only noise (overfitting), and it is possible that we retain just one source where in reality there are multiple relevant sources (underfitting).

Retaining a single source overfits the data if, in actuality, there is no source differentiating conditions at that frequency. This scenario can occur because the GED equations can be solved even for complete noise, essentially yielding sorted noise components. By assuming that the source with the largest eigenvalue meaningfully describes the data, we may mistake noise for a pattern. We mitigated potential overfitting by focusing on group-consistent peaks in the frequency spectra, relying on consistency across subjects to filter out noise-driven sources.

Conversely, retaining a single source underfits the data if multiple dissociable sources are active at that frequency. A single GED solution finds as many sources as there are sensors, more than one of which can be meaningful (de Cheveigné & Parra, 2014; Zuure et al., 2020). By retaining the largest source per frequency and discarding the rest, potentially valid sources are excluded from the model describing the data. We retained a single source per frequency regardless, on the theoretical grounds that the source with the largest eigenvalue is likely to be the most relevant, and on the practical grounds of convenience. Notably, this means that further interesting and relevant dynamics may exist that csGED is able to recover, but that we did not focus on.

By retaining a single source at each frequency in the spectrum, csGED may be subject to overfitting at some frequencies and underfitting at others. Future applications of csGED may simultaneously mitigate over- and underfitting through permutation testing. In this procedure, a null distribution of eigenvalues is constructed for each frequency, and sources with eigenvalues exceeding this null distribution are retained (e.g., Hayton et al., 2004; see Zuure et al., 2020, for a GED-specific example). Permutation testing may thus facilitate the detection of condition-relevant sources that do not occupy the largest eigenvalue.

### Pooling data across subjects

Like most source separation methods, csGED is a method to be applied to single subjects for which the results later have to be combined across subjects (cf. Parra et al., 2018). There are many possible approaches to pooling sources (e.g., clustering by topography, time series characteristics, or time-frequency characteristics; see also Bigdely-Shamlo et al., 2013; Huster et al., 2015; Huster & Raud, 2018; Onton et al., 2005; Pion-Tonachini et al., 2019). We here relied on the assumption that empirical sources occupying the largest eigenvalue and sharing a frequency are functionally similar. As the standard approach in electrophysiology is to aggregate data based purely on frequency band, our approach is consistent with standard practice. Other approaches may highlight additional consistencies across subjects. For example, sources may exist that share similar topographies and temporal characteristics between subjects, but that manifest at different frequencies.

### Extension beyond binary condition differences

csGED relies on the construction of a “signal” and a “reference” covariance matrix, meaning that the method lends itself to optimizing for a binary condition difference. One way to apply this method to multiple conditions is to compare all conditions pooled together against a baseline period; this would create a task-vs-baseline spatial filter that can be applied to the data from each individual condition. This approach also has the advantage of avoiding overfitting to a particular condition, albeit with a cost of reduced sensitivity to detecting subtle condition differences. csGED could also be extended to maximize a correlation with a variable over time or over trials, for example by integrating our frequency-specific spatial filters with the SPoC method (Dähne et al., 2014).

### Conclusions

csGED is a multivariate source separation technique that is optimized to reveal the frequency spectrum of binary condition differences. While demonstrated here in MEG data, it is a general method that is equally applicable to the analysis of multivariate neuroscience data at any spatial scale and with any measurement technique that provides repeated measurements. It carries the dual advantage of being hypothesis-driven in terms of conditions, which enhances its sensitivity to sources of interest, and being hypothesis-free in terms of frequency, which adds exploratory value. csGED thus has the potential to push the boundaries of our understanding in the analysis of cognitive electrophysiology.

## Acknowledgments

MBZ and MXC are funded by ERC-StG 638589. We thank Nadine Dijkstra for supplying the empirical data and providing insightful comments during analysis and writing.

## Supplementary material

**Supplementary Figure S1.**
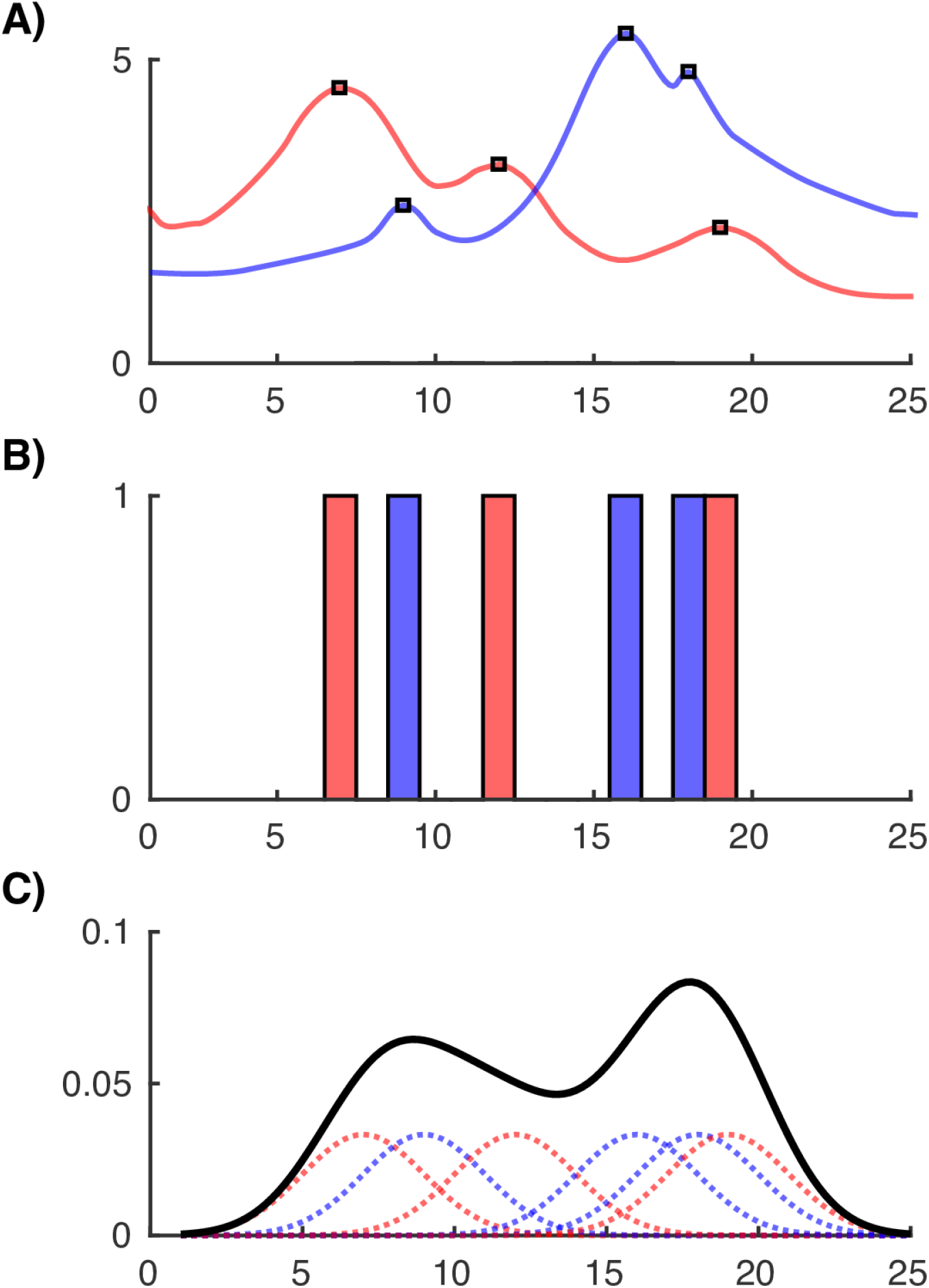
Illustration (using fabricated data) of the kernel density estimation procedure. **A)** The individual data (i.e., the eigenvalue spectra for each subject) to be transformed. **B)** The spectra are converted to a binary representation of the presence of eigenvalue peaks at each frequency. **C)** Gaussian distributions (dotted lines) that center on the individual peak frequencies are constructed. The sum of the Gaussian distributions, normalized by the number of peaks, is the KDF (black line).

**Supplementary Figure S2.**
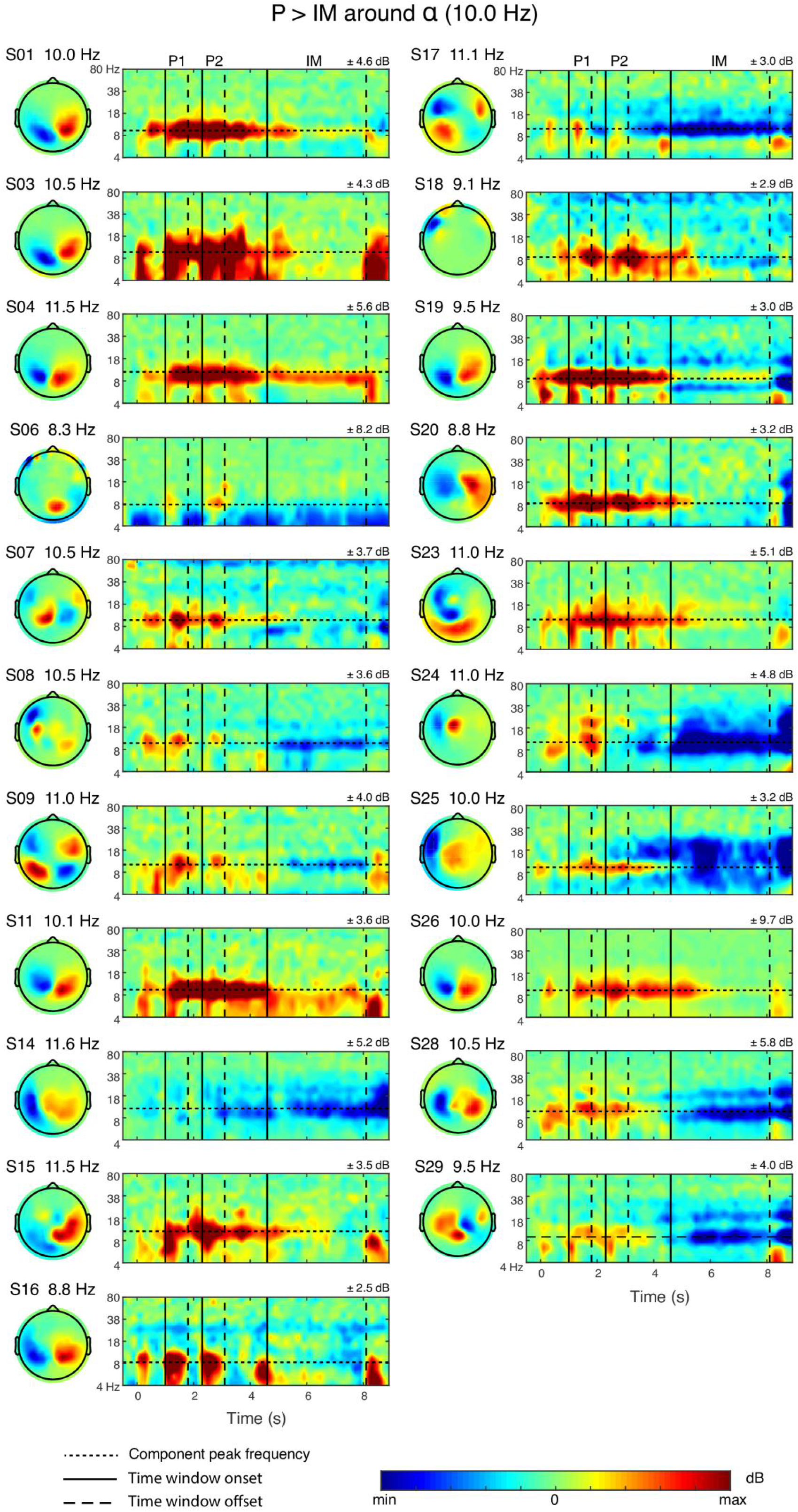
Individual subjects’ P > IM components around 10 Hz were diverse, but exhibited partial consistency in topography and time-frequency dynamics. Component spatial filters commonly captured an occipital topography. Components were characterized by alpha power increases during perception, alpha power decreases during imagery, or a combination of both. Horizontal dotted lines mark the filtering frequency at which each component was found. Vertical lines mark time window onset (solid) and offset (dashed), corresponding to the stimulus presentation for perception time windows (P1, P2) and the instructed imagery duration for the imagery time window (IM).

**Supplementary Figure S3.**
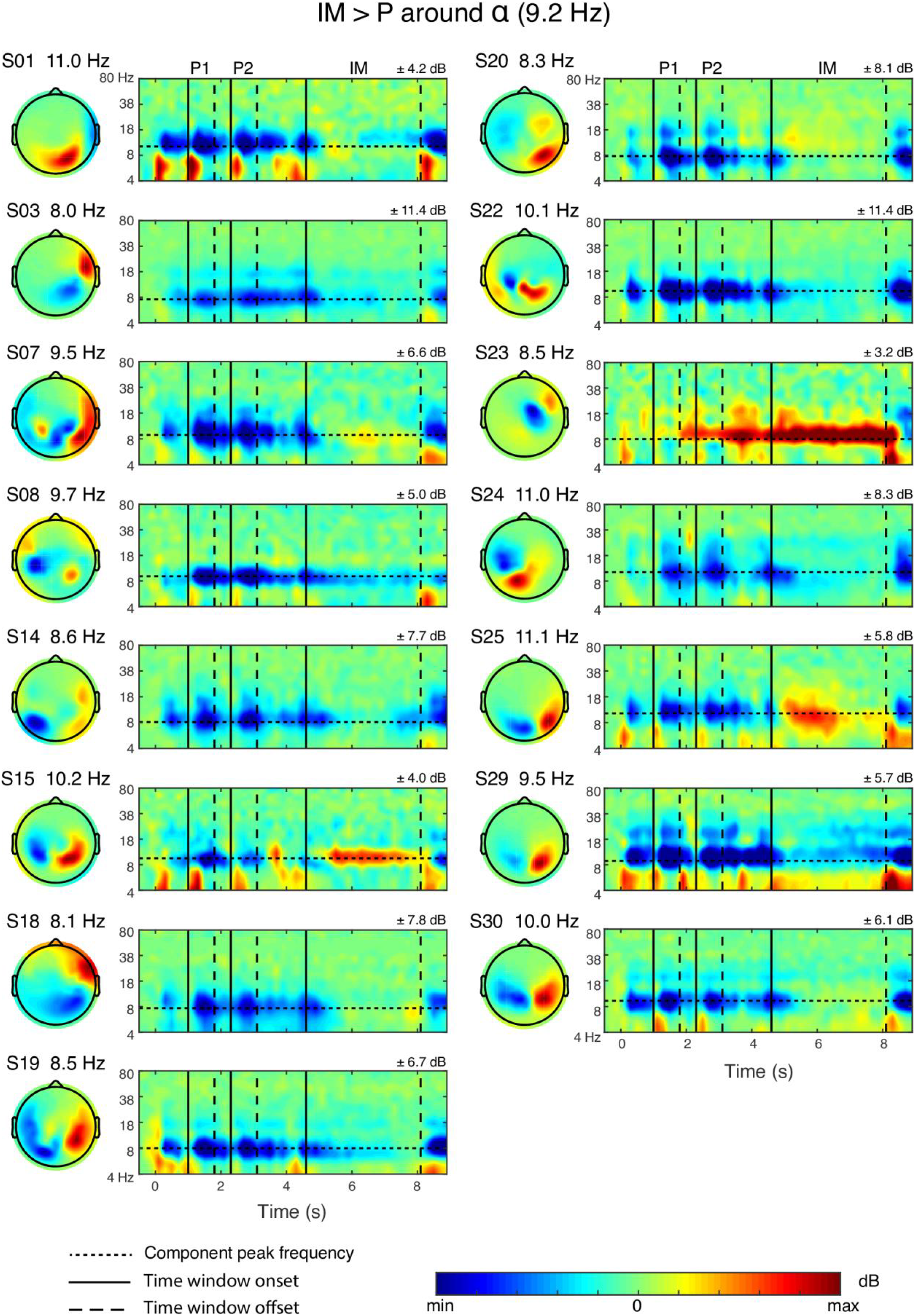
Same as Figure S2 but for the IM > P alpha peak. Component topographies were again commonly associated with an occipital topography. Components were characterized primarily by alpha power decreases during perception, and do not appear to be driven by imagery-related power changes.

**Supplementary Figure S4.**
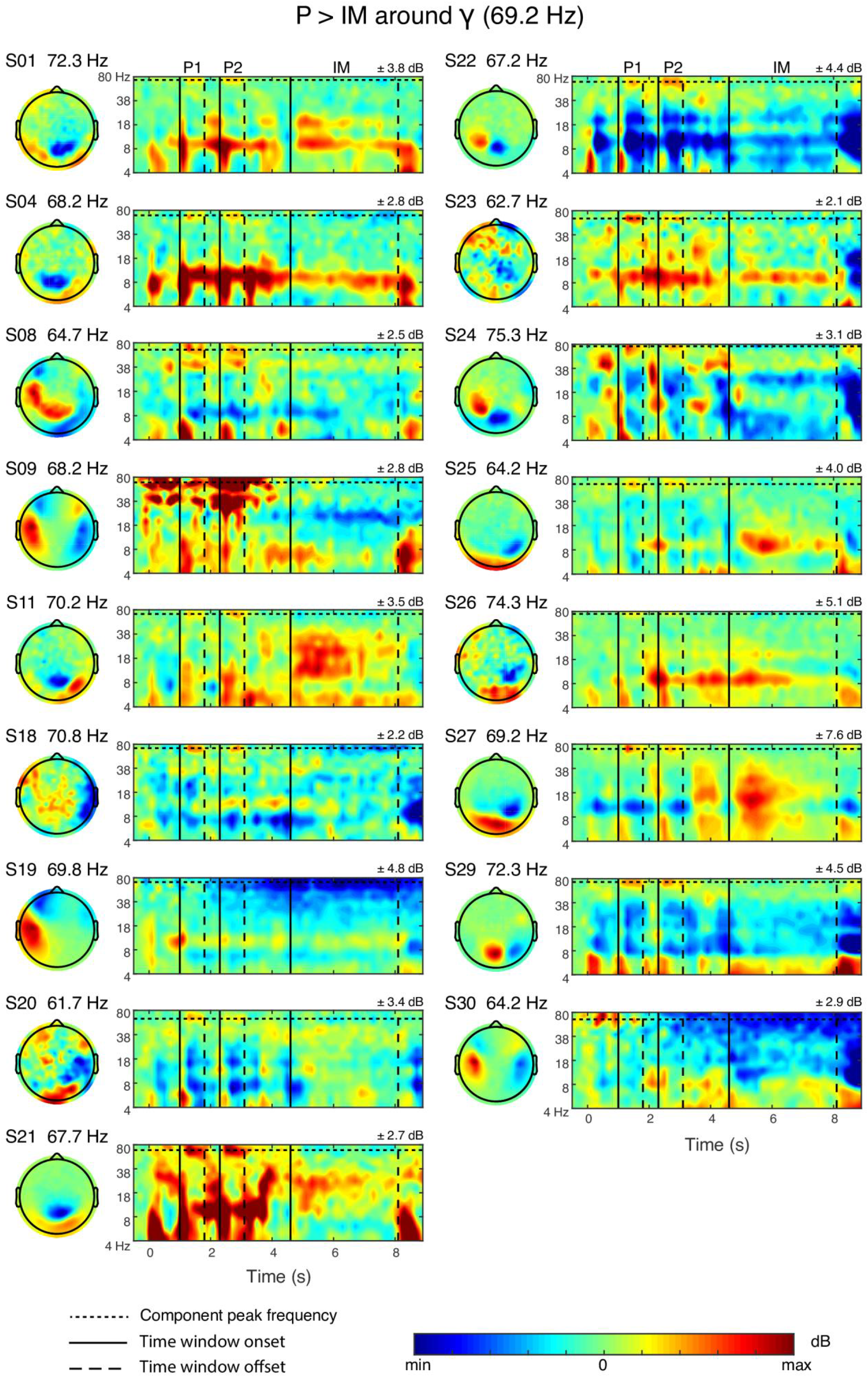
Same as Figure S2 but for the P > IM gamma peak. Components were characterized primarily by short-lived gamma power increases during perception.

**Supplementary Figure S5.**
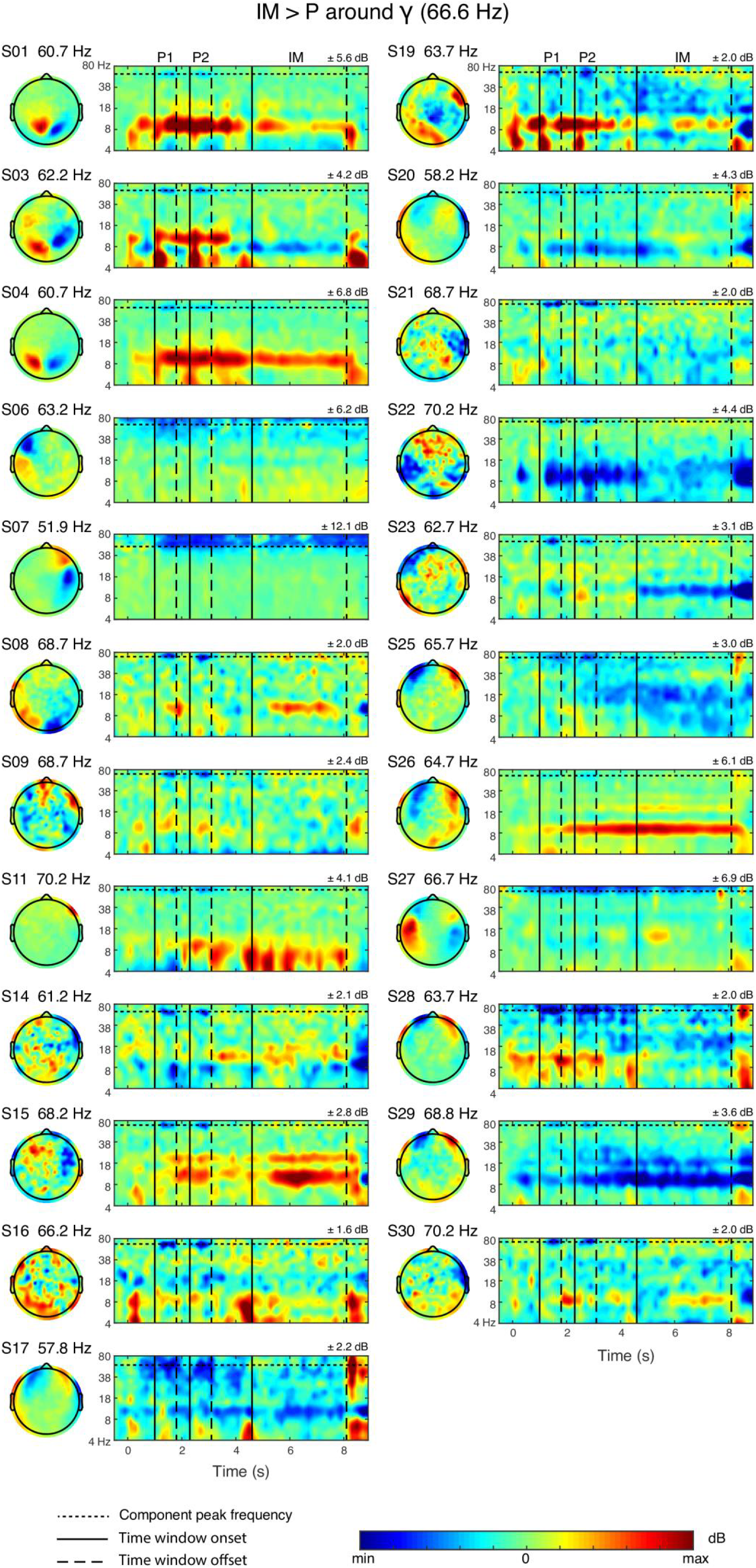
Same as Figure S2 but for the IM > P gamma peak. Components were characterized primarily by short-lived gamma power decreases during perception, and do not appear to be driven by imagery-related power changes.

## References

Asanowicz, D., Wołoszyn, K., Panek, B., & Wronka, E. (2019). On the locus of the effect of alerting on response conflict: An event-related EEG study with a speed-accuracy tradeoff manipulation. Biological Psychology, 145, 62–75.

Beckmann, C. F., & Smith, S. M. (2004). Probabilistic Independent Component Analysis for Functional Magnetic Resonance Imaging. In IEEE Transactions on Medical Imaging (Vol. 23, Issue 2, pp. 137–152). https://doi.org/10.1109/tmi.2003.822821

Bigdely-Shamlo, N., Mullen, T., Kreutz-Delgado, K., & Makeig, S. (2013). Measure projection analysis: a probabilistic approach to EEG source comparison and multi-subject inference. NeuroImage, 72, 287–303.

Brunton, B. W., Johnson, L. A., Ojemann, J. G., & Kutz, J. N. (2016). Extracting spatial-temporal coherent patterns in large-scale neural recordings using dynamic mode decomposition. Journal of Neuroscience Methods, 258, 1–15.

Cardoso, J. F., & Souloumiac, A. (1993). Blind beamforming for non-gaussian signals. In IEE Proceedings F Radar and Signal Processing (Vol. 140, Issue 6, p. 362). https://doi.org/10.1049/ip-f-2.1993.0054

Cohen, M. X. (2014). Analyzing Neural Time Series Data: Theory and Practice. The MIT Press.

Cohen, M. X. (2017a). Multivariate cross-frequency coupling via generalized eigendecomposition. eLife, 6. https://doi.org/10.7554/eLife.21792

Cohen, M. X. (2017b). Comparison of linear spatial filters for identifying oscillatory activity in multichannel data. Journal of Neuroscience Methods, 278, 1–12.

Cohen, M. X. (2020). A data-driven method to identify frequency boundaries in multichannel electrophysiology data. Journal of Neuroscience Methods, 347, 108949.

Dähne, S., Meinecke, F. C., Haufe, S., Höhne, J., Tangermann, M., Müller, K.-R., & Nikulin, V. V. (2014). SPoC: a novel framework for relating the amplitude of neuronal oscillations to behaviorally relevant parameters. NeuroImage, 86, 111–122.

de Cheveigné, A. (2019). ZapLine: a simple and effective method to remove power line artifacts. In bioRxiv (p. 782029). https://doi.org/10.1101/782029

de Cheveigné, A., & Arzounian, D. (2015). Scanning for oscillations. In Journal of Neural Engineering (Vol. 12, Issue 6, p. 066020). https://doi.org/10.1088/1741-2560/12/6/066020

de Cheveigné, A., & Parra, L. C. (2014). Joint decorrelation, a versatile tool for multichannel data analysis. NeuroImage, 98, 487–505.

Dijkstra, N., Bosch, S. E., & van Gerven, M. A. J. (2019). Shared Neural Mechanisms of Visual Perception and Imagery. Trends in Cognitive Sciences, 23(5), 423–434.

Dijkstra, N., Mostert, P., Lange, F. P. de, Bosch, S., & van Gerven, M. A. (2018). Differential temporal dynamics during visual imagery and perception. eLife, 7. https://doi.org/10.7554/eLife.33904

Fries, P., Womelsdorf, T., Oostenveld, R., & Desimone, R. (2008). The effects of visual stimulation and selective visual attention on rhythmic neuronal synchronization in macaque area V4. The Journal of Neuroscience: The Official Journal of the Society for Neuroscience, 28(18), 4823–4835.

Haufe, S., Meinecke, F., Görgen, K., Dähne, S., Haynes, J.-D., Blankertz, B., & Bießmann, F. (2014). On the interpretation of weight vectors of linear models in multivariate neuroimaging. NeuroImage, 87, 96–110.

Hayton, J. C., Allen, D. G., & Scarpello, V. (2004). Factor Retention Decisions in Exploratory Factor Analysis: a Tutorial on Parallel Analysis. In Organizational Research Methods (Vol. 7, Issue 2, pp. 191–205). https://doi.org/10.1177/1094428104263675

Hoogenboom, N., Schoffelen, J.-M., Oostenveld, R., Parkes, L. M., & Fries, P. (2006). Localizing human visual gamma-band activity in frequency, time and space. NeuroImage, 29(3), 764–773.

Huster, R. J., Plis, S. M., & Calhoun, V. D. (2015). Group-level component analyses of EEG:validation and evaluation. Frontiers in Neuroscience, 9, 254.

Huster, R. J., & Raud, L. (2018). A Tutorial Review on Multi-subject Decomposition of EEG. Brain Topography, 31(1), 3–16.

Klimesch, W. (2018). The frequency architecture of brain and brain body oscillations: an analysis. The European Journal of Neuroscience, 48(7), 2431–2453.

Klimesch, W., Sauseng, P., & Hanslmayr, S. (2007). EEG alpha oscillations: The inhibition– timing hypothesis. In Brain Research Reviews (Vol. 53, Issue 1, pp. 63–88). https://doi.org/10.1016/j.brainresrev.2006.06.003

Nikulin, V. V., Nolte, G., & Curio, G. (2011). A novel method for reliable and fast extraction of neuronal EEG/MEG oscillations on the basis of spatio-spectral decomposition. NeuroImage, 55(4), 1528–1535.

Nunez, P. L., & Srinivasan, R. (2006). Electric Fields of the Brain. https://doi.org/10.1093/acprof:oso/9780195050387.001.0001

Onton, J., Delorme, A., & Makeig, S. (2005). Frontal midline EEG dynamics during working memory. NeuroImage, 27(2), 341–356.

Oostenveld, R., Fries, P., Maris, E., & Schoffelen, J.-M. (2011). FieldTrip: Open source software for advanced analysis of MEG, EEG, and invasive electrophysiological data. Computational Intelligence and Neuroscience, 2011, 156869.

Parra, L. C., Haufe, S., & Dmochowski, J. P. (2018). Correlated Components Analysis - Extracting Reliable Dimensions in Multivariate Data. In arXiv [stat.ML]. arXiv. http://arxiv.org/abs/1801.08881

Parra, L. C., Spence, C. D., Gerson, A. D., & Sajda, P. (2005). Recipes for the linear analysis of EEG. NeuroImage, 28(2), 326–341.

Pion-Tonachini, L., Kreutz-Delgado, K., & Makeig, S. (2019). ICLabel: An automated electroencephalographic independent component classifier, dataset, and website. NeuroImage, 198, 181–197.

Sedley, W., & Cunningham, M. O. (2013). Do cortical gamma oscillations promote or suppress perception? An under-asked question with an over-assumed answer. In Frontiers in Human Neuroscience (Vol. 7). https://doi.org/10.3389/fnhum.2013.00595

Tadel, F., Baillet, S., Mosher, J. C., Pantazis, D., & Leahy, R. M. (2011). Brainstorm: a user-friendly application for MEG/EEG analysis. Computational Intelligence and Neuroscience, 2011, 879716.

Thut, G., Nietzel, A., Brandt, S. A., & Pascual-Leone, A. (2006). Alpha-band electroencephalographic activity over occipital cortex indexes visuospatial attention bias and predicts visual target detection. The Journal of Neuroscience: The Official Journal of the Society for Neuroscience, 26(37), 9494–9502.

Tiedt, H. O., Lueschow, A., Pauls, A., & Weber, J. E. (2016). The face-responsive M170 is modulated by sensor selection: An example of circularity in the analysis of MEG-data. Journal of Neuroscience Methods, 266, 137–140.

Vidaurre, D., Myers, N. E., Stokes, M., Nobre, A. C., & Woolrich, M. W. (2019). Temporally Unconstrained Decoding Reveals Consistent but Time-Varying Stages of Stimulus Processing. Cerebral Cortex, 29(2), 863–874.

Wang, Y., & Guo, Y. (2019). A hierarchical independent component analysis model for longitudinal neuroimaging studies. NeuroImage, 189, 380–400.

Yang, J., Zhang, D., Frangi, A. F., & Yang, J.-Y. (2004). Two-dimensional pca: a new approach to appearance-based face representation and recognition. In IEEE Transactions on Pattern Analysis and Machine Intelligence (Vol. 26, Issue 1, pp. 131–137). https://doi.org/10.1109/tpami.2004.1261097

Zuure, M. B., Hinkley, L. B., Tiesinga, P. H. E., Nagarajan, S. S., & Cohen, M. X. (2020). Multiple Midfrontal Thetas Revealed by Source Separation of Simultaneous MEG and EEG. The Journal of Neuroscience: The Official Journal of the Society for Neuroscience, 40(40), 7702–7713.

