## Supplementary figures and images for "Narrowband multivariate source separation for semi-blind discovery of experiment contrasts"

### Supplementary Figure S1

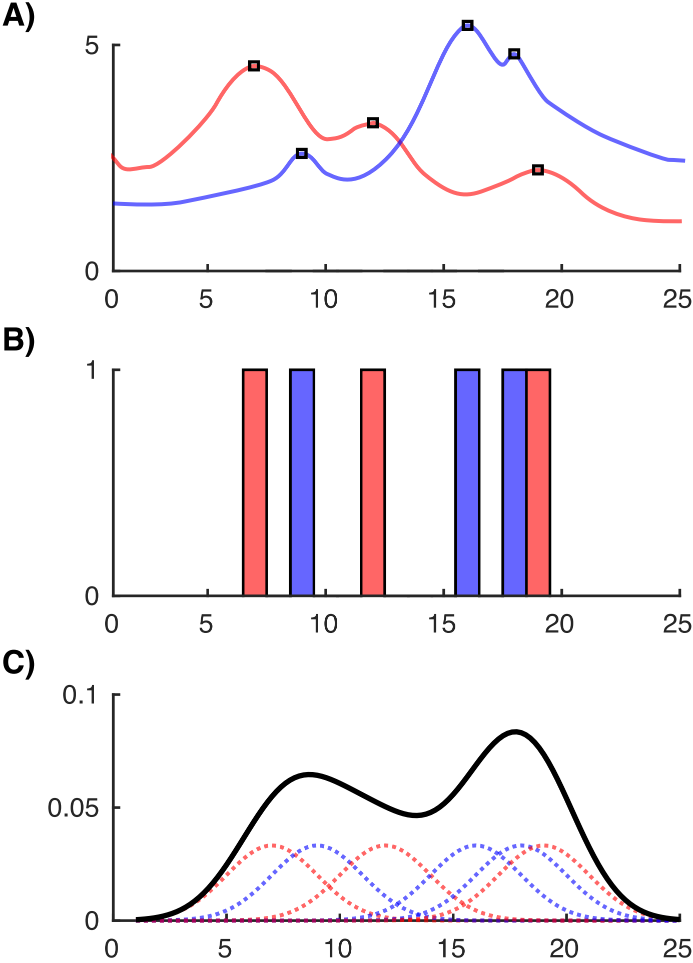

### Supplementary Figure S2

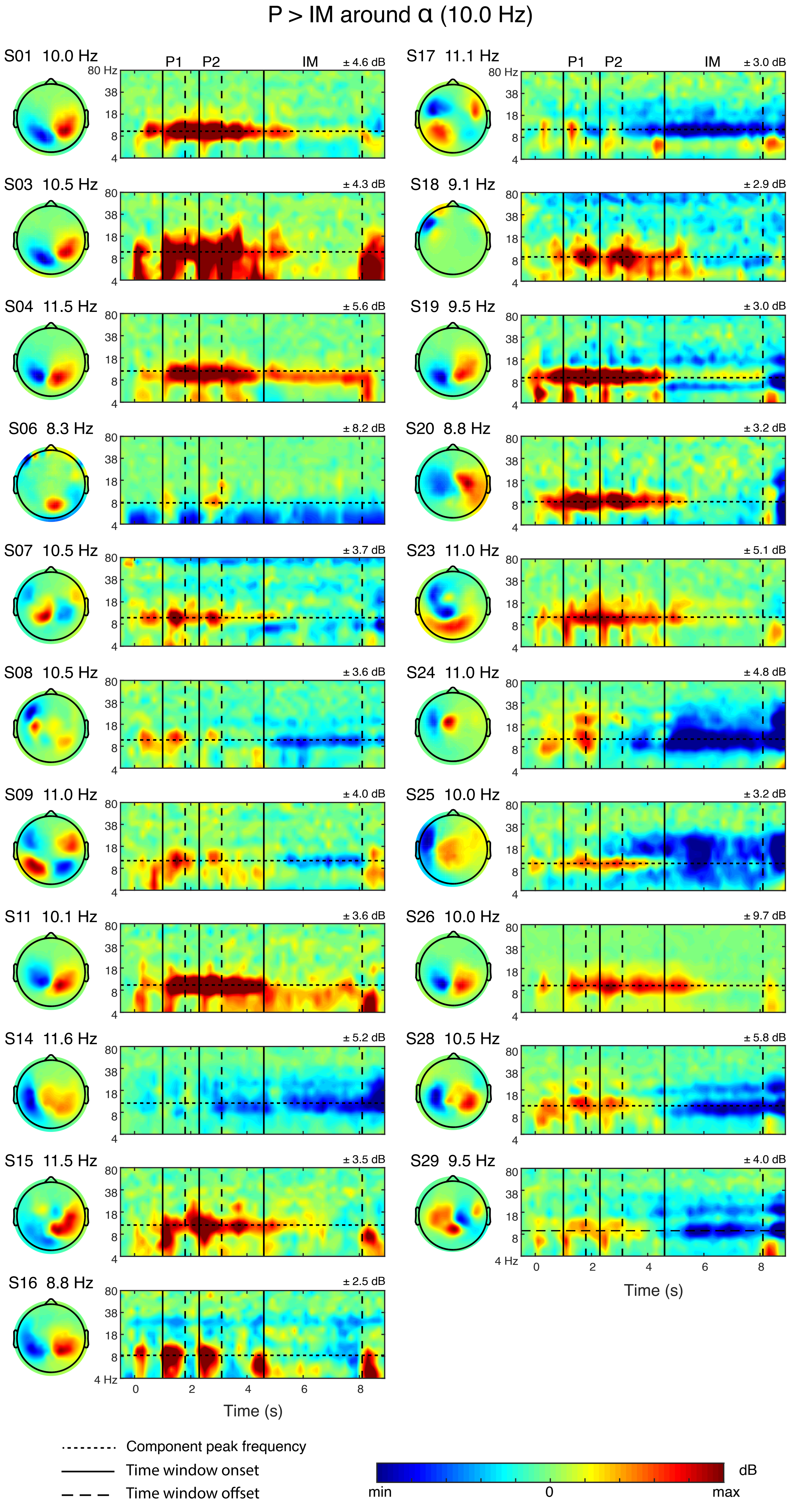

### Supplementary Figure S3

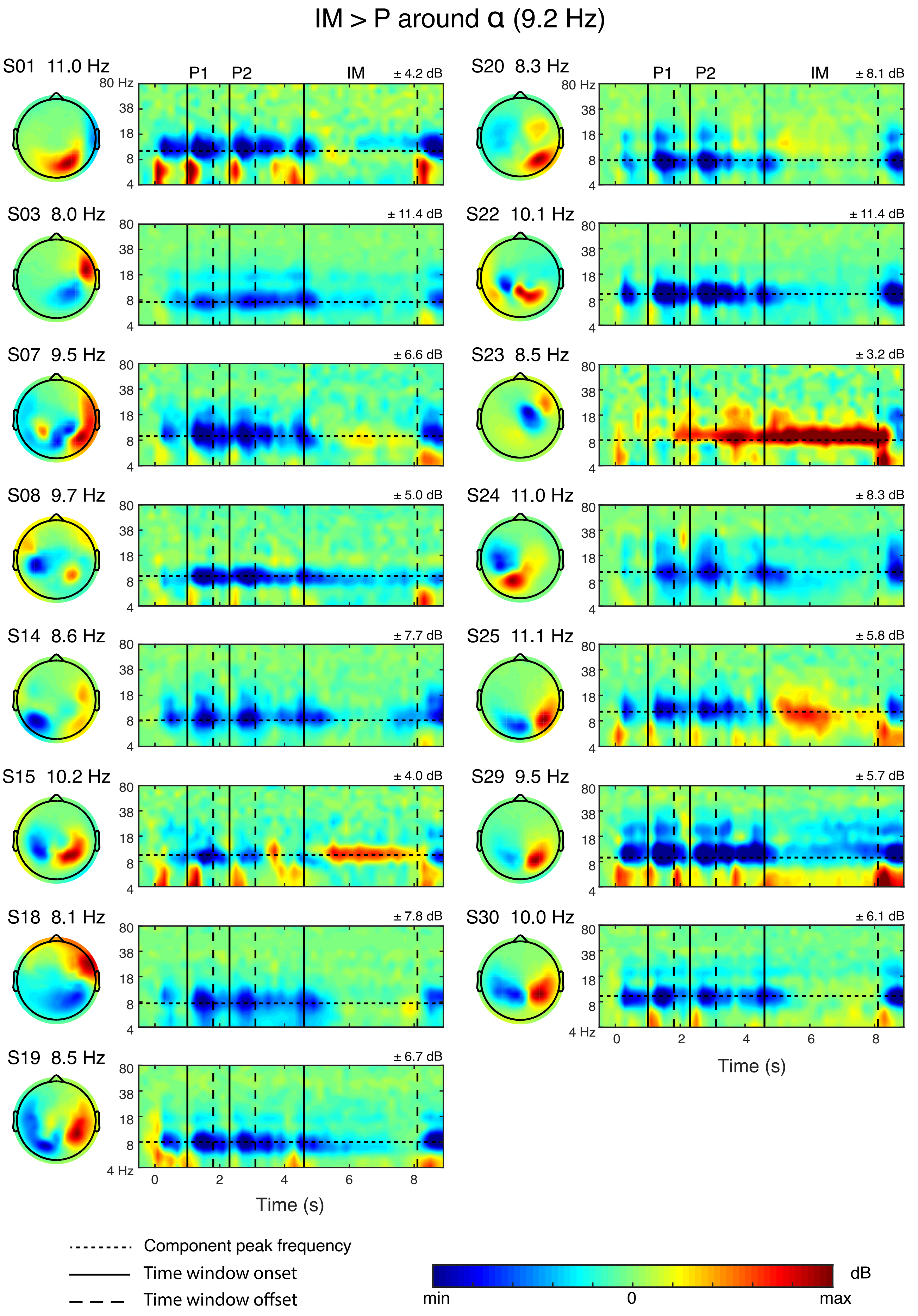

### Supplementary Figure S4

#### P > IM around γ (69.2 Hz)

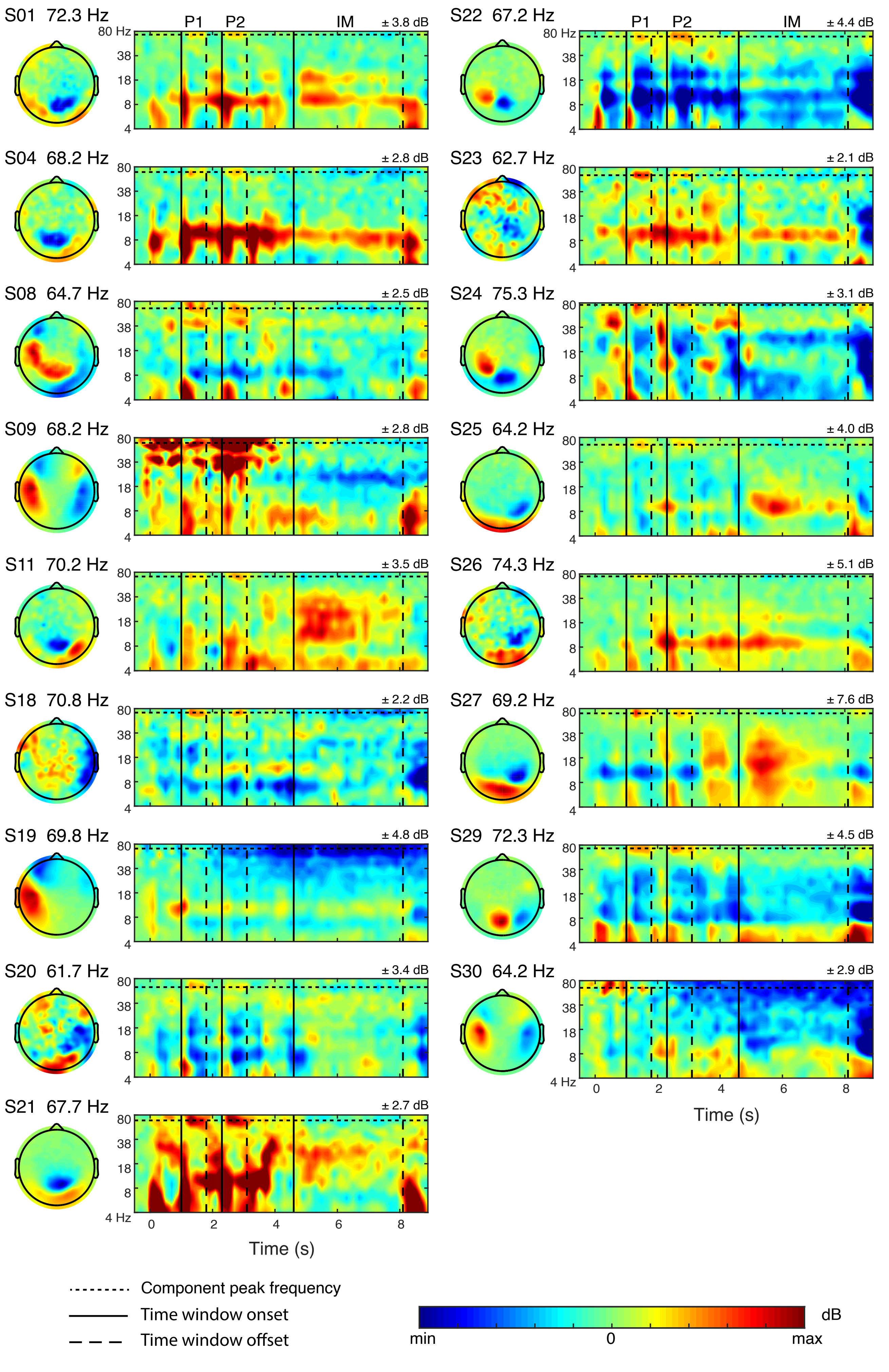

### Supplementary Figure S5

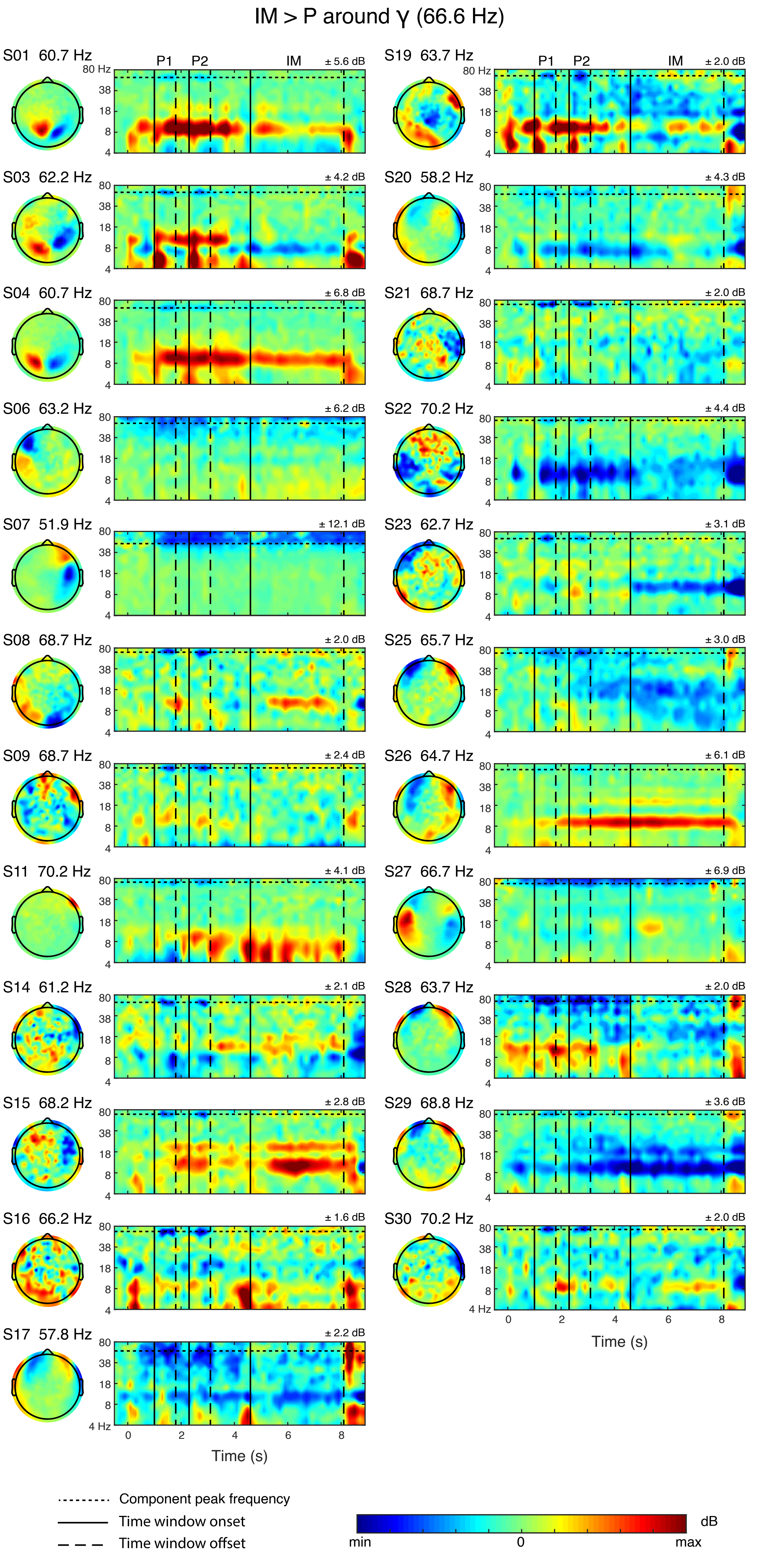
